# A comparison of the binding sites of antibodies and single-domain antibodies

**DOI:** 10.1101/2023.05.30.542890

**Authors:** Gemma L. Gordon, Henriette L. Capel, Bora Guloglu, Eve Richardson, Ryan L. Stafford, Charlotte M. Deane

## Abstract

Antibodies are the largest class of biotherapeutics. However, in recent years, single-domain antibodies have gained traction due to their smaller size and comparable binding affinity. Antibodies (Abs) and single-domain antibodies (sdAbs) differ in the structures of their binding sites: most significantly, single-domain antibodies lack a light chain and so have just three CDR loops. Given this inherent structural difference, it is important to understand whether Abs and sdAbs are distinguishable in how they engage a binding partner and thus, whether they are suited to different types of epitopes. In this study, we use non-redundant sequence and structural datasets to compare the paratopes, epitopes and antigen interactions of Abs and sdAbs. We demonstrate that even though sd-Abs have smaller paratopes, they target epitopes of equal size to those targeted by Abs. To achieve this, the paratopes of sdAbs contribute more interactions per residue than the paratopes of Abs. Additionally, we find that conserved framework residues are of increased importance in the paratopes of sd-Abs, suggesting that they include non-specific interactions to achieve comparable affinity. Furthermore, the epitopes of sdAbs and Abs cannot be distinguished by their shape. For our datasets, sd-Abs do not target more concave epitopes than Abs: we posit that this may be explained by differences in the orientation and compaction of sdAb and Ab CDR-H3 loops. Overall, our results have important implications for the engineering and humanization of sdAbs, as well as the selection of the best modality for targeting a particular epitope.

## Introduction

Monoclonal antibodies are widely used as biotherapeutics, but their high molecular weight (∼ 150kDa) can cause high production costs as well as poor diffusion rates that limit tissue penetration (*1*–*3*). These properties of antibodies (Abs) have led to increased interest in recent years around smaller antibody fragments such as single-domain antibodies (sdAbs). SdAbs are isolated VH domains (VHHs) homologous to the VH domain in antibodies and are derived primarily from camelid heavy-chain antibodies (*4*). SdAbs are approximately one tenth the mass of antibodies (∼ 15kDa). Despite significantly reduced available diversity, sdAbs have been shown to achieve comparable binding specificity and affinities to Abs (*5*). Furthermore, sdAbs are thermostable and have shown higher solubility, blood clearance and tissue penetration than Abs (*2, 6, 7*). These properties suggest that sdAbs have huge potential in therapeutic use, provided they can be successfully humanized (*8*).

Major structural differences exist between sdAbs and Abs, the most conspicuous being that sdAbs lack a light chain and therefore have only three complementarity-determining region (CDR) loops, half that of Abs. The CDR loops in both Abs and sdAbs are known to contain the majority of the binding site. Understanding the differences in the binding sites of these two classes of immunoglobulin, in terms of how their structures enable interaction with their binding partners, would facilitate decision-making as to which modality might be more effective when targeting a particular epitope.

In previous work, Zavrtanik *et al*. (2018) (*9*), suggested that sdAbs target more “rigid, concave, conserved and structured” epitopes. This hypothesis that sdAbs target more concave epitopes is often linked to the fact that the CDR-H3 loops of sdAbs are longer than those of conventional Abs (*10, 11*). Zavrtanik *et al*. (2018) and Mitchell and Colwell (2018a) (*12*) found an average difference in loop length of between three and four residues. Many papers have theorised that the longer CDR-H3 loops of sdAbs can protrude into concave spaces in a protein antigen surface that would be inaccessible to a conventional Ab with a shorter CDR-H3 loop (*13*–*16*). However, as highlighted by Henry and Mackenzie (2018) (*17*), isolated case studies make up much of the supporting literature on this idea: whether the epitope space of sdAbs is truly more concave than that of Abs has not yet been fully assessed.

Aside from differences in CDR-H3 loop length, previous comparisons of the paratopes of sdAbs and Abs have shown that sdAbs have more hydrophobic character than Abs but are similarly enriched in aromatic residues (*9*). Furthermore, sdAbs tend to draw more residues from framework regions into the paratope, whereas Abs are more reliant on the CDR loops to interact with an antigen (Ag) (*9, 12*).

The previous studies of Zavrtanik *et al*. (*9*) and Mitchell and Colwell (*12, 18*) are limited by their relatively small datasets: Zavrtanik *et al*. analyse 105 sdAb-Ag crystal complexes, while Mitchell and Colwell compare sets of 90 sdAb-Ag and Ab-Ag crystal complexes (2018a) and then 156 sdAb-Ag and Ab-Ag complexes (2018b).

As sdAb datasets have increased in size in recent years (*19*), we have examined the binding sites of sdAbs and Abs using non-redundant datasets of 892 Ab-Ag and 345 sdAb-Ag structural complexes, alongside non-redundant datasets of 1,614,526 human VH sequences (from Eliyahu *et al*., 2018 (*20*)) and 1,596,446 camel VHH sequences (from Li *et al*., 2016 (*21*)). We find that in agreement with previous work, the paratopes of sd-Abs are smaller, on average, than those of Abs and that the CDR-H3 loop of sdAbs is longer. In our analysis, the paratopes of sdAbs and Abs cannot be distinguished by amino acid composition. We also find that the epitopes of sdAbs and Abs cannot be differentiated by their size, shape, or amino acid composition, contrasting with previous suggestions that the epitopes of sdAbs are more concave than those of Abs. Overall, our results suggest that sdAbs and Abs do not target different epitopes despite differences in their paratopes. However, they may be distinguishable by the manner in which they interact with these epitopes. We find that a greater number of interactions per residue are initiated by the CDR-H3 loop of sdAbs and that the framework region of sdAbs contributes more residues to the paratope. These differences likely contribute to the ability of sdAbs to achieve comparable binding affinity to Abs. However, our analysis shows that many of the binding framework residues are conserved positions, suggesting that sdAb binding may include non-specific interactions.

## Methods

### Sequence datasets

Non-redundant sequence datasets were obtained from the Observed Antibody Space (OAS) database (*22*). A set of 1,621,889 human VH sequences generated by Eliyahu *et al*. (2018) (*20*) and 1,601,636 camel VHH sequences generated by Li *et al*. (2016) (*21*), were filtered to remove duplicated sequences. Final datasets, referred to as the “Abs sequence dataset” and “sdAbs sequence dataset”, consist of 1,614,526 human VH sequences and 1,596,446 camel VHH sequences. These sequence datasets were used to compare the CDR lengths and the amino acid compositions of framework residues and CDR loops between Abs and sdAbs.

### Structure datasets

We created up-to-date, non-redundant datasets of both Abs and sdAbs that were in complex with protein antigens (Ags). We refer to these as the “Abs structural dataset” and “sdAbs structural dataset”. These structures were extracted from SAbDab (*23*) and SAbDab-nano (*19*) on the 23^rd^ February 2022. The datasets were extracted as follows:

1. Only Ab-Ag and sdAb-Ag complexes for which at least one of the CDR residues of the antibody is in close contact, defined as under 4.5Å, to the antigen.
2. Only the Abs and sdAbs identified as in a complex with a protein antigen (> 50 residues), according to SAbDab annotations.
3. Only structures of complexes solved by X-ray crystallography to *≤* 3.0Å resolution.
4. Abs and sdAbs were filtered separately to remove redundancy using a sequence identity cut-off of 95% across the IMGT-defined CDR residues using CD-HIT (*24*).
5. A small number of complexes were reintroduced if their epitope identity score was less than 75% compared to any other complex, to include complexes containing similar CDRs but different epitopes. To calculate epitope identity, epitope sequences were first aligned using CD-HIT (*24*). Based on the aligned positions, the epitope identity score was determined as the fraction of matching (distance-defined) epitope residues (same amino acids and same aligned position) across the epitope residues of the two antigens.

The resulting sdAbs structural dataset consisted of 345 complexes, of which 309 had “unique” CDRs. The final Abs structural dataset consisted of 892 complexes, of which 792 had “unique” CDRs. SI Text S1 and Table S1 give further detail on dataset curation and a breakdown of the number of complexes remaining at each filtering step. Table S2 shows species variation for both structural datasets. Supplementary Figure S1 shows distributions of epitope identity across datasets.

### Numbering definitions

The IMGT numbering scheme and CDR definitions were used throughout this work (CDR1: IMGT residues 27-38, CDR2: IMGT residues 56-65, CDR3: IMGT residues 105-117 (*25*). ANARCI (*26*) was used to number all of the Abs and sdAbs.

### Binding site definitions

We describe the binding site using three definitions. As used in most methods annotating and predicting paratopes or epitopes, we consider a distance definition, which includes all antibody residues which are in close contact with the antigen (≤ 4.5Å). A very similar result is achieved by defining the binding site by solvent-accessible surface area (SASA), where residues are included in the paratope or epitope if they become buried on complex formation (SASA-defined). In our work we focus on defining the binding site by the interactions occurring between pairs of residues, using Arpeggio (*27*). Arpeggio determines interaction types based on distance, angle, and atom type. It was run on each PDB file in both structure datasets after cleaning with the associated cleaning script (*https://github.com/harryjubb/pdbtools*), using a distance threshold of 4.5Å. This generates a five-bit fingerprint for each pairwise interatomic contact which shows the type of interactions occurring. These include, van der Waals, steric clashes, covalent bonds, proximal interactions (defined as being within the cut-off distance but not representing a meaningful interaction) and specific interactions such as hydrogen bonds. This output was processed to exclude interactions with water molecules and chains other than the antibody and antigen. Heterogens were removed with BioPython (*28*). For all remaining positions, the interatomic interactions were summarised per residue-residue pair. Residues were considered to interact if at least one of the atom-atom pairs in these residues established a van der Waals (vdW) bond or a specific interaction. Clashing vdW and proximal interactions were classified as contacts if no specific bonds were observed. We refer to this latter definition of the binding site as the interactions-defined paratope and interactions-defined epitope.

Interatomic interactions between the Ab:Ag and sdAb:Ag complexes were compared by counting the total number observed. If multiple interaction types were identified between a single pair of atoms, the interactions were counted individually. Mean and standard deviation of the observed interactions were calculated by sub-sampling 10% of the total set of interactions 1000 times.

Supplementary Figures S2 and S3 visualise the difference between paratopes and epitopes defined by contacts or interactions, and the difference between each definition of the binding site.

### Amino acid composition

The sequence datasets were used to compare compositions of CDR loops. The sdAbs and Abs sequence datasets were split by germline and only those belonging to IGHV3 compared: this included all sequences for the sdAbs dataset but reduced the Abs dataset to 761,235 sequences. Sequences were aligned using ANARCI numbering annotation. The proportions of individual amino acids at each position in each CDR-H loop were determined. Positions were omitted where less than 5% of sequences had an amino acid at that position.

To assess the conservation of framework residues that appear in the paratope, firstly the structural datasets were used to determine which positions are often involved in the paratopes of sdAbs and Abs. Framework residues were considered as important contributors to the paratope if they were observed in at least 10% of the complexes in our datasets. The amino acid compositions of these same positions were then obtained from the sequence datasets as a background for comparison.

### Epitope shape

To determine the concavity of the epitope, protein structures were represented as three-dimensional alpha shapes. Alpha shapes are a generalisation of the convex hull, derived from Delaunay triangulation, and can be used to characterise the geometry of a surface, such as that of a protein (*29, 30*). The Python packages open3d v0.15.1 (*31*) and trimesh v3.12.9 (2019) (*32*) were used to generate a mesh from a point cloud of PDB atomic coordinates and then calculate vertex defect values for all vertices of the mesh.

Vertex defects values indicate how concave or convex the epitope surface is: more positive values correspond to more convex regions; more negative values correspond to more concave regions. As the atomic coordinates of the PDB structure and the vertex coordinates of the mesh do not map perfectly, a K-d tree was generated with scipy v1.7.2 (*33*) to determine which vertex coordinates were closest (based on Euclidean distance) to the atomic coordinates, allowing the vertex defects at the epitope surface to be extracted.

The accuracy of the mesh representation was validated by aligning the epitope regions of the mesh with the PDB structure and determining the root-mean-square deviation (RMSD) with the Kabsch algorithm (*34*). Alpha parameters when generating a mesh were set for each complex in the range of 0-3, such that the resulting RMSD between the mesh and PDB coordinates of the epitope was *≤* 1Å.

### Canonical forms of the CDRs

Canonical forms of sdAb and Ab structures were identified using the PyIgClassify2 database (*35*).

### Structural clustering

Antibody chains from the 345 sdAb:Ag and 892 Ab:Ag complexes were extracted, giving 301 and 838 unique sdAbs and Abs structures (as some PDB entries include sdAbs or Abs that form complexes with multiple antigens). A greedy clustering method was used where each of the sets of CDR-H1, CDR-H2 and CDR-H3 loops were clustered based on their length and RMSD with a cut-off of 1.5Å. The number of clusters which contain both sdAbs and Abs (overlap clusters) was determined. The expected number of overlap clusters was found by generating random clusters of matching size. Random clusters were generated 20 times from the original set of all Ab and sdAb structures and the mean and standard deviations for the number of overlap clusters was calculated.

### Orientation of CDR-H3 loops

We analysed the general orientation of the CDR-H3 loops of Abs and sdAbs by examining their centers of geometry in reference to an R3 coordinate system (see Text S2 for method and Supplementary Figure S4). The dataset used for this analysis includes the structures of 388 bound sdAbs, 116 unbound sdAbs, 1977 bound Abs and 862 unbound Abs. Structures were downloaded from SAbDab (*23*) on 8^th^ August 2022 and generated individually to be non-redundant at 95% sequence identity. Structures were numbered with the IMGT scheme using ANARCI (*26*) and CDR definitions used accordingly. Any structures with missing backbone atoms in CDR-H loops or anchors (three residues on either side of each loop) were also removed.

Using the spherical coordinates method, *ρ* describes the reach of the CDR-H3 loop away from the rest of the VH domain. A CDR-H3 loop in an extended conformation will have a high *ρ* value whereas a loop of identical length that is folded against the VH domain will have a lower value. *ϕ* gives an indication of whether the CDR-H3 loop is horizontally oriented towards the rest of the VH domain or away from it. In the case of Ab structures, a high *ϕ* value indicates packing against the VL domain. *θ* gives a measure of the elevation of the loop. A low value corresponds to a CDR-H3 that extends directly up and away from the rest of the VH domain, whereas a high value indicates that the loop is “folding” down. In the case of Ab structures, a high *θ* value corresponds to a loop that is packed into the groove created by the VH-VL interface. Lastly, we divide the loop length by *ρ* to give a measure of compaction. A loop with low compactness uses its entire length to reach away from the VH domain, whereas high compactness corresponds to a loop that is packed against the VH.

### Statistical tests

As not all distributions followed the normal distribution, significant differences between the sdAbs and Abs were tested by bootstrap re-sampling in which 5000 bootstrap samples are taken of size 300. The unpaired mean difference and the p-value of the two-sided permutation t-test are reported. Results are described as significant for p-value < 0.05.

### Visualisations

All visualisations were created using open-source PyMOL v2.4.1 (*36*), UCSF ChimeraX (*37*), or matplotlib v3.5.1 (*38*).

### Code and data availability

Code written for the analyses, alongside structural and sequence datasets are available at *github*.*com/oxpig*.

## Results

In this study, non-redundant sequence datasets for Abs and sdAbs of size 1,614,526 and 1,596,446 respectively, and non-redundant structural datasets of 892 Ab-Ag and 345 sdAb-Ag complexes, were compared with respect to their paratopes, epitopes and their interactions with their respective antigens to identify the differences and similarities between their binding sites, and to determine whether these two modalities target different types of epitopes.

### The CDR-H3 loop is longer in sdAbs than in Abs

Previous work has shown that the CDR-H3 loops of sd-Abs are longer than those of Abs. Lengths of the CDR loops were compared for both sequence and structural datasets. When comparing the sdAbs and Abs sequence datasets, we find that the CDR-H1 loops of Abs are, on average, slightly longer than those of sdAbs by 0.4 residues. Abs have on average longer CDR-H2 loops by 0.2 residues. The CDR-H3 loops are significantly longer in sdAbs by 1.4 residues on average (Figure 1A). The results from the structural dataset are consistent with the trends observed for the sequence datasets: for the solved structures, bootstrap re-sampling shows that for CDR-H1, there is a significant difference between sdAbs and Abs of 0.2 residues and for CDR-H2, we find that there is a difference of 0.08 (p-value = 0.12). For the structural datasets, the CDR-H3 loop is significantly longer in sdAbs than in Abs by 1.6 residues (Figure 1B). This finding agrees with previous studies.

**Figure 1.**
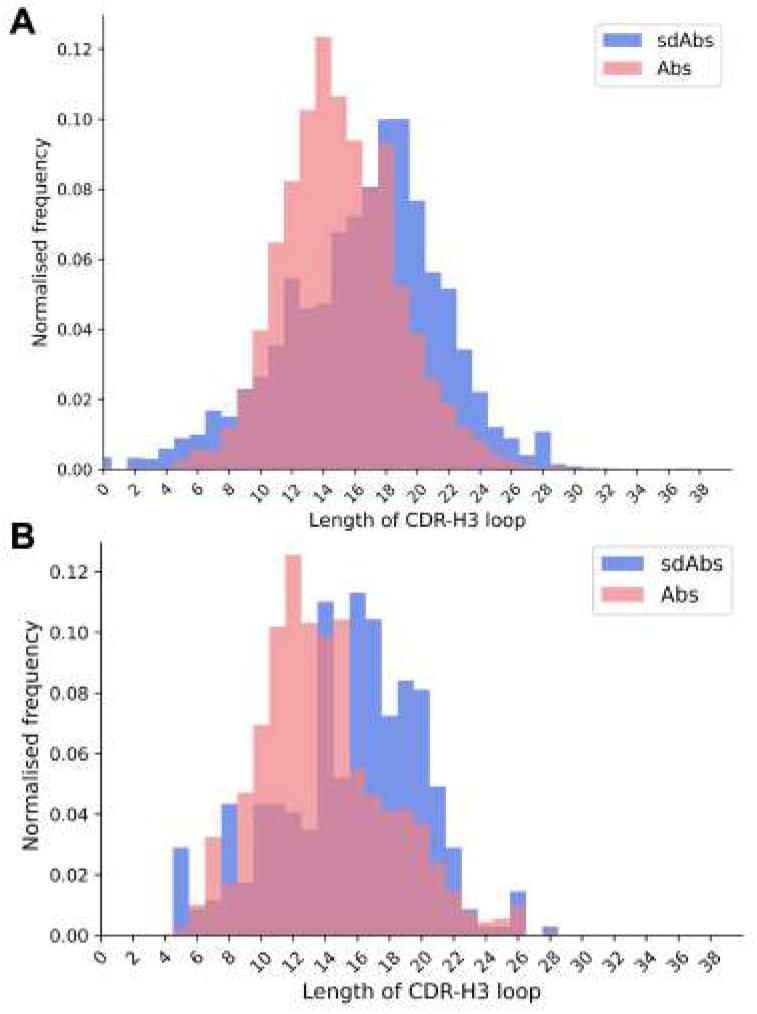
The distributions of CDR-H3 loop length for (**A**) sequence data and (**B**) structural data both show that CDR-H3 loops in sdAbs (blue) tend to be longer than those in Abs (pink).

### Structural clustering shows a separation between Abs and sdAbs CDR structures

Further to comparing the lengths of the CDR loops found in Abs and sdAbs, we next structurally clustered the CDR loops to determine whether they adopt distinct conformations and occupy different regions of structural space. If Abs and sdAbs were to adopt different paratope shapes, this would suggest that the epitopes they are able to bind would differ.

Our initial approach was to assign canonical forms to each of the Abs and sdAbs loop structures, according to updated canonical forms from Kelow *et al*. (2022) (*35*).

However, for both Abs and sdAbs a significant percentage of CDR loops could not be assigned a canonical form. Therefore, CDR loops were clustered based on length and RMSD, with a cut-off of 1.5Å. Clustering of the CDR loops of our 838 Abs and 301 sdAbs structures collectively returned 168 clusters for CDR-H1, 94 clusters for CDR-H2 and, as expected given the differences in CDR-H3 length and the high variability of CDR-H3 in general, 729 CDR-H3 clusters.

The number of clusters containing both Abs and sdAbs structures was determined and a mean and standard deviation for the expected number of overlap clusters, if random clustering had occurred, was calculated (Table 1). For CDR-H1, 18 clusters contained both Abs and sd-Abs compared to an expected value of 16.2 ± 1.29 for random clusters. For CDR-H1, 23 clusters contained both Abs and sdAbs compared to an expected value of 22.3 ± 0.829. Overall, we observe that for all CDR-H loops, the number of clusters we see with both Abs and sdAbs occurring within them is within the range of what would be expected had the structures been clustered at random. This indicates that sdAbs and Abs may assume distinct CDR conformations. As the CDR loops form the majority of the binding site, this suggests that Abs and sdAbs may prefer to bind in different ways.

**Table 1.** Clustering the structures of the CDR loops of sdAbs and Abs based on length and conformational similarity (measured by RMSD) shows that there is overlap between the shapes that CDR loops of sdAbs and Abs form. However, this overlap is within the range of that observed on random clustering, and as such suggests that sdAbs and Abs adopt distinct CDR conformations. Values in the table show the number of structures within each cluster, with the number of clusters containing only a single structure shown in brackets.

|  | CDR loop |  |  |
| --- | --- | --- | --- |
|  | CDR-H1 | CDR-H2 | CDR-H3 |
| **Abs-only**<br>**(single-occupancy)** | 66 (48) | 35 (21) | 489 (383) |
| **SdAbs-only**<br>**(single-occupancy)** | 84 (64) | 36 (25) | 230 (197) |
| **Overlap** | 18 | 23 | 10 |
| **Total** | 168 | 94 | 729 |

### SdAbs and Abs cannot be distinguished by the amino acid composition of their CDR loops

We next examined the CDR loop sequences belonging to IGHV3 germlines, taken from the sdAbs and Abs sequence datasets. This reduced the size of the Abs dataset to 761,235 sequences (all 1,596,446 sequences in the sdAbs sequence dataset belong to the IGHV3 germline). Sequences within each dataset were aligned via ANARCI annotation and the amino acid composition at each position in each loop determined. Positions were omitted if less than 5% of sequences in a dataset had a residue at that position. Supplementary Figure S5 shows sequence logo plots of the CDR loops of Abs and sdAbs.

Given the size of the sequence space, the probability of finding the same sequences in both Abs and sd-Abs CDR loops is low. The expected proportion of identical sequences between the sdAbs and Abs sequences for each loop was calculated and compared to the actual overlap. For CDR-H1, the expected overlap is 6.31×10^−11^ versus 0.024, for CDR-H2, 7.33 × 10^−11^ versus 0.021, and for H3, 1.53 × 10^−21^ versus 3.00 × 10^−4^. As the actual number of identical sequences is greater than the expected number, this suggests that there are similarities in the amino acid compositions of sdAbs and Abs CDR loops, which likely arise from their similar genetic background.

### SdAbs paratopes are significantly smaller than those of Abs

In addition to assessing differences in the CDR loops of Abs and sdAbs, we considered whether there are overall differences in their respective paratopes by comparing their size. Previous work has revealed that sd-Abs can show comparable binding affinity to Abs despite their smaller size (*5*). Given that sdAbs are missing the VL domain and therefore half of an Ab potential binding site, we would expect them to also have a smaller paratope. Using our non-redundant structural datasets, we compared the size of sdAb and Ab paratopes for each of the distance-defined, interactions-defined and SASA-defined paratopes. We found that for distance-defined paratopes, sdAb paratopes are significantly smaller than Ab paratopes by 3.6 residues and for interaction-defined paratopes, SdAb paratopes are smaller than Ab paratopes by 2.6 residues (Figure 2). Supplementary Figure S6 shows results consistent with the above for the SASA-defined paratopes. The differences found between the CDRs and more specifically the paratopes of sdAbs and Abs in our datasets suggest that these two modalities may target distinct epitopes.

**Figure 2.**
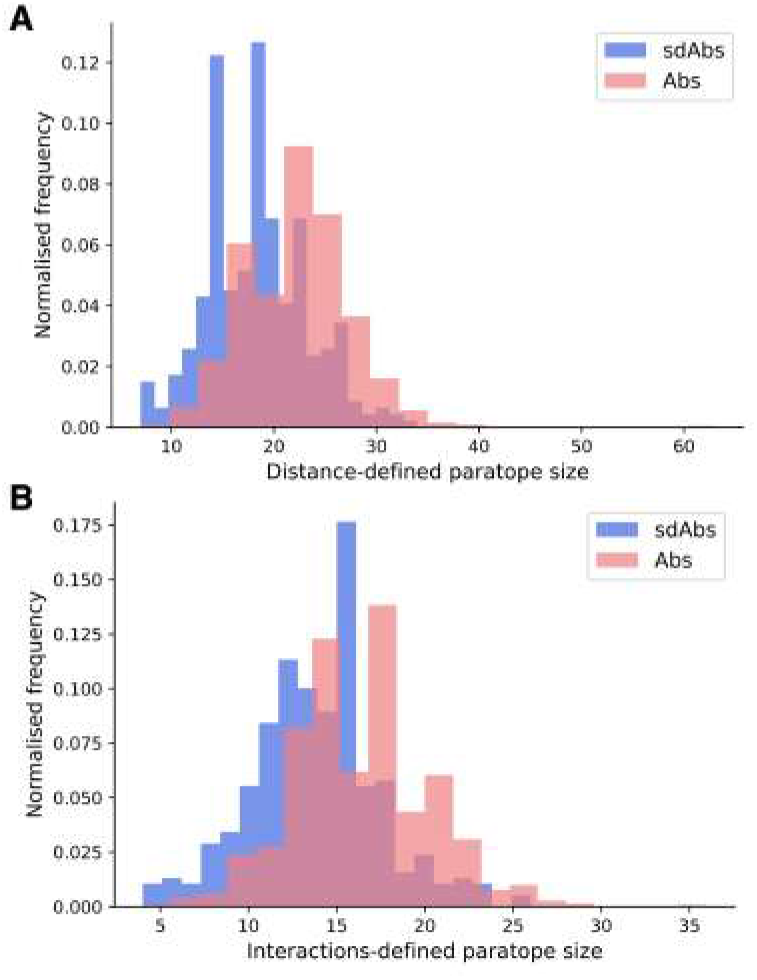
The paratopes of sdAbs (blue) tend to be smaller than the paratopes of Abs (pink). **(A)** Distributions of the sizes of the distance-defined paratopes, where sdAbs paratopes are significantly smaller by 3.6 residues on average compared to Abs. **(B)** Distributions of the sizes of the interactions-defined paratopes, where sdAbs paratopes are significantly smaller by 2.6 residues, on average.

### Epitopes targeted by sdAbs and Abs have similar amino acid compositions

We next assessed the epitopes of Abs and sdAbs. One factor that may differ between sdAbs and Abs is the amino acid compositions of their target epitopes. Due to the relatively small size of the structural datasets, following the work of Wong *et al*., 2021 (*39*), amino acid compositions for the epitopes were determined by classifying amino acids into seven classes (aliphatic, aromatic, sulfur, hydroxyl, basic, acidic and amine). For each epitope, the fraction of each observed class was determined and the distributions of amino acid types for epitopes of sdAbs and Abs were compared.

Comparisons of the seven classes for both distance-defined and interactions-defined epitopes reveal that for epitopes of sdAbs, there is a small but significant increase in the number of aromatic residues, and a significant decrease in the number of basic residues (Supplementary Figures S7, S8). These results suggest small differences, but based on our relatively small structural datasets, it is hard to conclude that the epitopes of sd-Abs and Abs are distinct in terms of amino acid composition.

### Epitopes of Abs are more linear than those of sdAbs

Epitopes are often characterised by whether they are more linear or discontinuous in nature. A linear epitope is formed from amino acid residues that fall next to each other at the primary sequence level, whereas a discontinuous epitope is formed from residues that are not adjacent in the amino acid sequence but are pulled together upon folding (*40, 41*). Here, we determined whether Abs and sdAbs show distinct epitope preferences in terms of epitope continuity. We represent how continuous an epitope is by the number of contiguous residues in the epitope sequence.

For both the distance and interactions-based definitions, epitopes of Abs tend to be slightly more linear than those of sdAbs (Figure 3). Abs showed a significantly greater percentage of linear residues for both the distance-defined (4.6%) and interactions-defined (6.9%) epitopes. Similar results are observed when comparing the raw count of linear residues (Supplementary Figure S9). Results are replicated for the SASA-defined epitopes (Supplementary Figure S10). As the epitopes of sdAbs and Abs are of comparable size, the fact that Abs have slightly more linear epitopes than sd-Abs is not due to a difference in epitope size.

**Figure 3.**
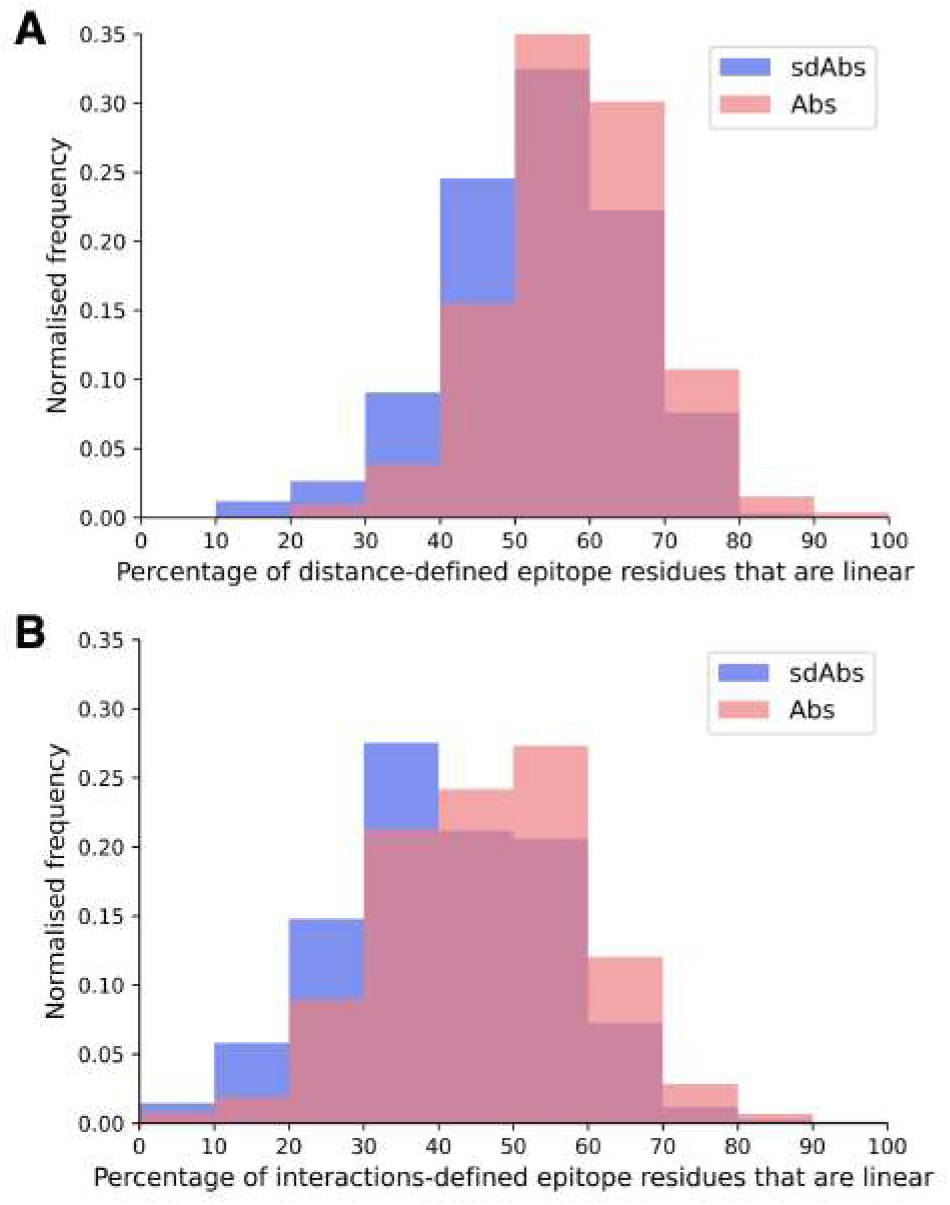
The epitopes targeted by Abs are relatively more linear than epitopes targeted by sdAbs, as suggested by the distributions of percentages of linear residues for epitopes targeted by Abs (pink) and sdAbs (blue) for the **(A)** distance-defined epitopes and **(B)** interactions-defined epitopes.

### Epitopes targeted by sdAbs and Abs are of comparable size

As described above, the paratopes of sdAbs are smaller than those of Abs, which suggests that sdAbs may be limited to binding smaller epitopes. Here, we determined the number of residues in the distance-defined epitopes, the SASA-defined epitopes and the interactions-defined epitopes for our non-redundant structural datasets. Our results show that for each of our epitope definitions, there is no significant difference between the size of the epitopes targeted by sdAbs and Abs (Figure 4, Supplementary Figure S11). Despite their smaller paratope size, sdAbs target epitopes of equal size to those targeted by Abs. This indicates that the paratopes of sdAbs must interact with their epitopes in a different way to that of Abs paratopes.

**Figure 4.**
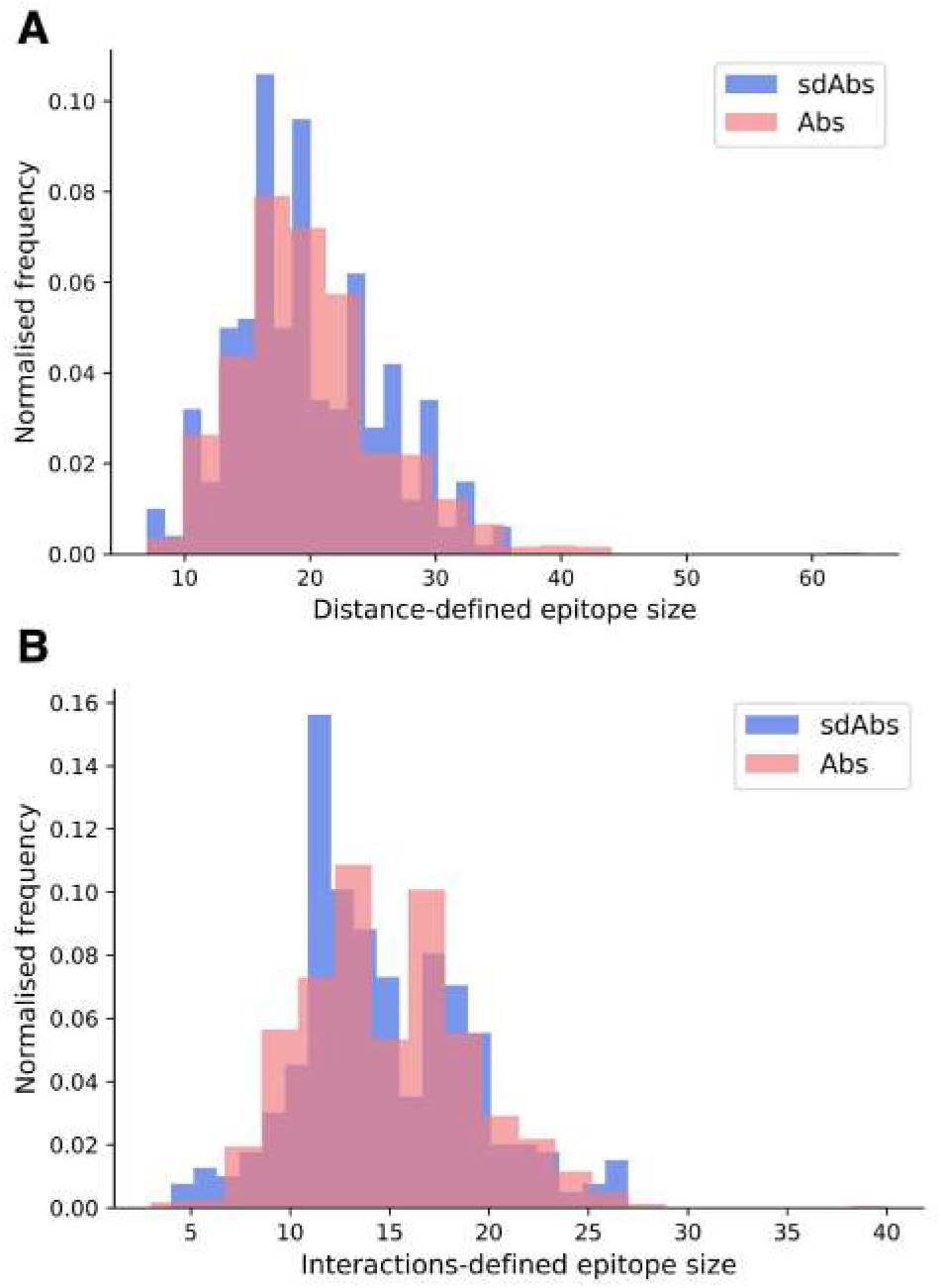
SdAbs are able to target epitopes of equal size to those targeted by conventional Abs, as suggested by the distributions of the number of residues in the **(A)** distance-defined epitopes for Abs (pink) and sdAbs (blue) structural datasets, where a mean difference of 0.59 is observed between sdAbs and Abs (p-value = 0.22) and **(B)** interaction-defined epitopes, where a mean difference of 0.32 is observed between sdAbs and Abs (p-value = 0.34).

### Epitopes targeted by sdAbs are not more concave than those of Abs

In agreement with existing studies on smaller datasets, we found that sdAbs have longer CDR-H3 loops than Abs. Previous work has suggested that this facilitates interactions between sdAbs and more concave epitopes compared to Abs (*5, 11, 13*–*16*). To assess whether the epitopes of sdAbs do indeed tend to be more concave, the surface shapes of all interaction-defined epitopes of sdAbs and Abs were analysed.

An alpha shape representation of the antigen surface was used. To ensure this was an accurate representation of the crystal structure, for each structure the mesh representation vertex coordinates were aligned to the PDB atomic coordinates. The RMSD values of all alignments were determined and plotted against the corresponding mean vertex defect value for each epitope. There is no correlation between the RMSD and mean vertex defects value for the epitope, allowing confidence in the accuracy of the surface representation(Supplementary Figure S12). The most convex and most concave epitopes (based on mean vertex defects (VD) value) for both sdAbs and Abs are shown in Supplementary Figure S13.

The VD values were determined for the mesh representation of each protein antigen surface, and the VD values correlating with the epitope region were found. The VD value indicates the curvature of the surface at a particular vertex. An area with many positive VD values will be a more convex region and similarly, an area with many negative values will be more concave. The mean of all VD values of the epitope, and the proportion of negative VD values across the epitope, were used to represent the curvature of each epitope.

The unpaired mean difference between sdAbs and Abs mean VD values was 0.076 (p-value = 0.13) (Figure 5). The unpaired mean difference between sdAbs and Abs in the proportions of negative VD values, which indicate a concave shape, was -0.0089 (p-value = 0.15) (Supplementary Figure S14). These results suggest that there is no significant difference between the concavity of epitopes targeted by sdAbs and Abs, contradicting previous studies (*9*). Furthermore, the similarity in epitope shapes we observe does not appear to be due more recently released structures changing the distributions of concavity: for our structural datasets, the similarity between the shapes of Abs and sdAbs epitopes has not markedly changed over time (Supplementary Figure S15).

**Figure 5.**
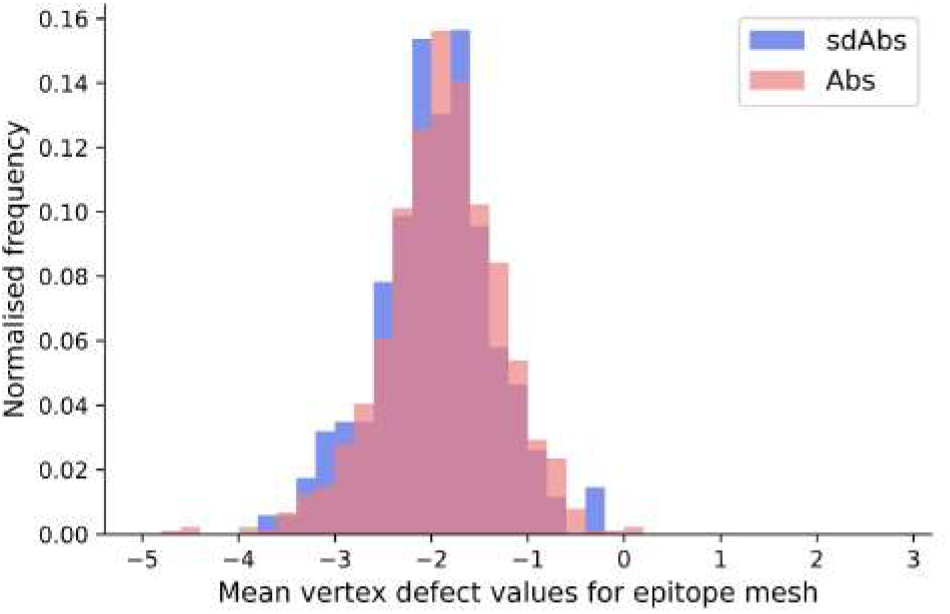
SdAbs, in general, do not target more concave epitopes than Abs. Distributions of mean vertex defects values for the interactions-defined epitopes of sdAbs (blue) and Abs (pink). The mean vertex defects value represents the concavity of the epitope, where more a positive value correlates with a more convex epitope, and a more negative value with a more concave epitope. The unpaired mean difference between sdAbs and Abs mean vertex defects values was 0.076 (p-value = 0.13).

### CDR-H3 loop length does not correlate with epitope concavity

The hypothesis that sdAbs are generally able to target more concave epitopes derives from the finding that their CDR-H3 loops are longer than those of Abs (*10, 11*). However there is no correlation between the length of the CDR-H3 loop and the curvature of the epitope surface for our datasets (Figure 6). For sdAbs, the Pearson correlation coefficient for mean VD values against the CDR-H3 loop length was -0.016, whilst the coefficient for proportion of negative VD values for an epitope against the CDR-H3 loop length was 0.039. For Abs, the Pearson correlation coefficient for mean VD values against the CDR-H3 loop length was -0.063, whilst the coefficient for the proportion of negative VD values for an epitope against the CDR-H3 loop length was 0.030. These results indicate that the length of the CDR-H3 loop alone does not influence the shape of the epitope targeted by either antibody type.

**Figure 6.**
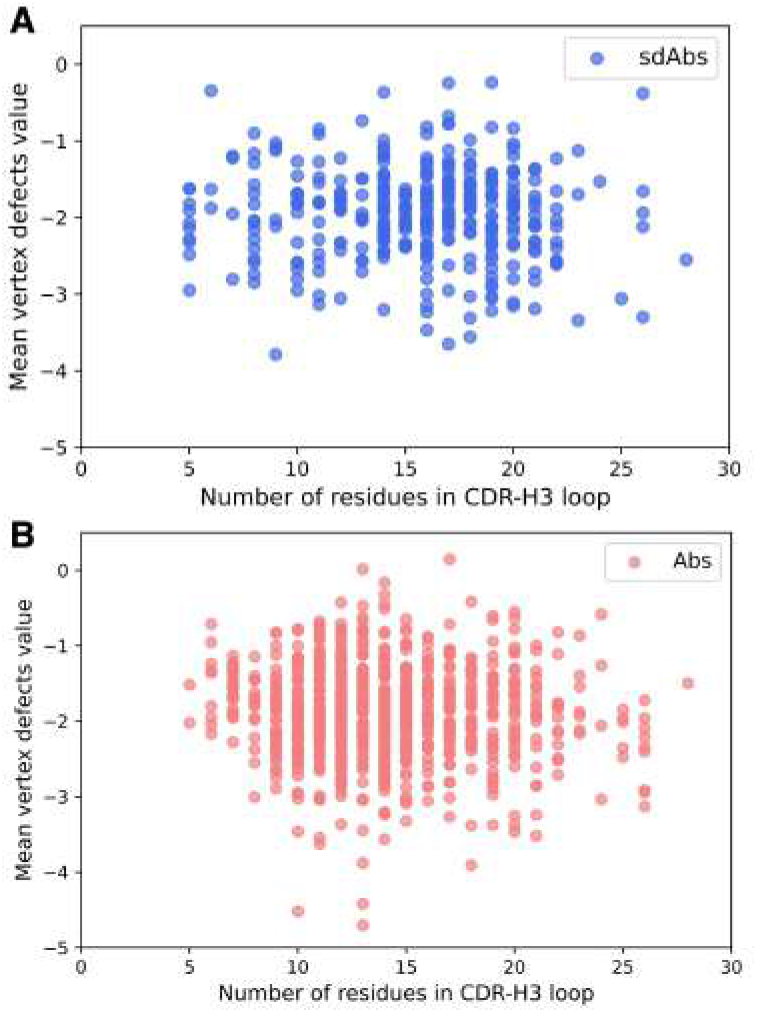
There is no correlation between the length of the CDR-H3 loop and the concavity of the epitope structure for either Abs or sdAbs. **(A)** Correlation between mean vertex defects values of sdAb epitopes and length of CDR-H3 loop. **(B)** Correlation between mean vertex defects values of Ab epitopes and length of CDR-H3 loop.

### Abs and sdAbs target epitopes of similar shape due to packing of sdAb CDR-H3 loops against the VHH domain

In light of our finding that the length of the CDR-H3 loop does not dictate the shape of the epitope to which a paratope binds, we examined the differences in the orientation of Ab and sdAb CDR-H3 loops relative to the rest of the VH domain, to determine how the conformation of the CDR-H3 loop may affect epitope preference.

We use four descriptors to describe the orientation of the CDR-H3 loops (see Methods, Supplementary Text S2 and Supplementary Figure S4): the parameter *ρ* represents the reach of the CDR-H3 loop away from the VH domain, *ϕ* describes the horizontal orientation of the CDR-H3 towards the rest of the VHH (for sdAbs), or against the VL domain (for Abs), *θ* describes loop extension where a low value corresponds to a CDR-H3 extending up and away from the rest of the VH domain and lastly compaction, which is determined by dividing loop length by *ρ*.

Near-identical distributions of *ρ* values suggests that the two types of antibodies have similar reach, indicating that sdAbs cannot necessarily provide extended paratopes via their CDR-H3 loops compared to Abs (Figure 7A). A shoulder in the distribution of *ρ* values for Abs above the median value suggests that Abs may be more able to target deeper epitopes that require a longer reach.

**Figure 7.**
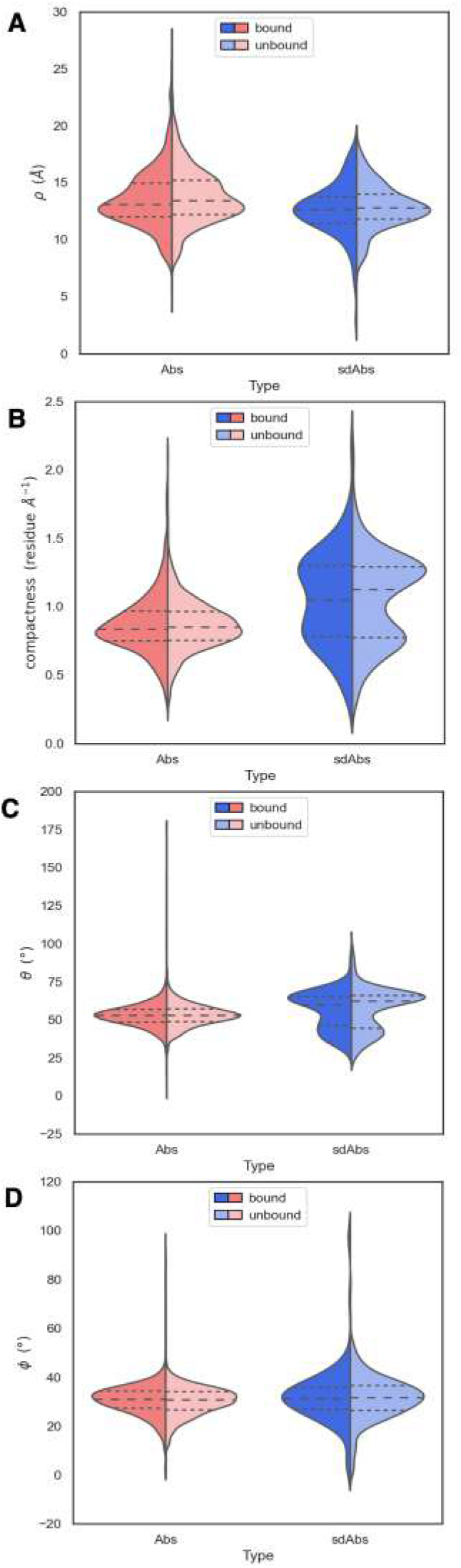
The orientation of the CDR-H3 loops of sdAbs (blue) suggests why sdAbs do not target more concave epitopes than Abs (pink). **(A)** Distributions of *ρ* values show that sdAbs and Abs have similar reach. **(B)** On average, sdAb CDR-H3 loops are more compacted than Ab loops. **(C)** Distributions of *θ* values indicate that the majority of sdAb CDR-H3 loops do not extend upwards away from the VHH domain, but lie flat against it. **(D)** Distributions of *ϕ* values indicate that the majority of sdAb CDR-H3 loops pack against the VHH domain. In all figures the bound examples are shown in a darker shade on the left of the distributions, with the unbound in a lighter shade on the right.

The observation that sdAb CDR-H3 loops tend to be longer than those in Abs, whilst having similar reach, may be explained by loop compaction. On average, sdAb CDR-H3 loops are much more compacted than Ab loops (Figure 7B). The distribution of compactness scores for sdAbs is bimodal, with the first peak corresponding to the distribution found in Abs. This suggests one population of sdAb CDR-H3 loops that behaves similarly to Ab CDR-H3 loops, and one population that is more folded against the VHH domain (Figure 8A). SdAbs can either increase their reach with CDR-H3 length at a rate similar to Abs, or their loops can remain in a more heavily compacted state.

**Figure 8.**
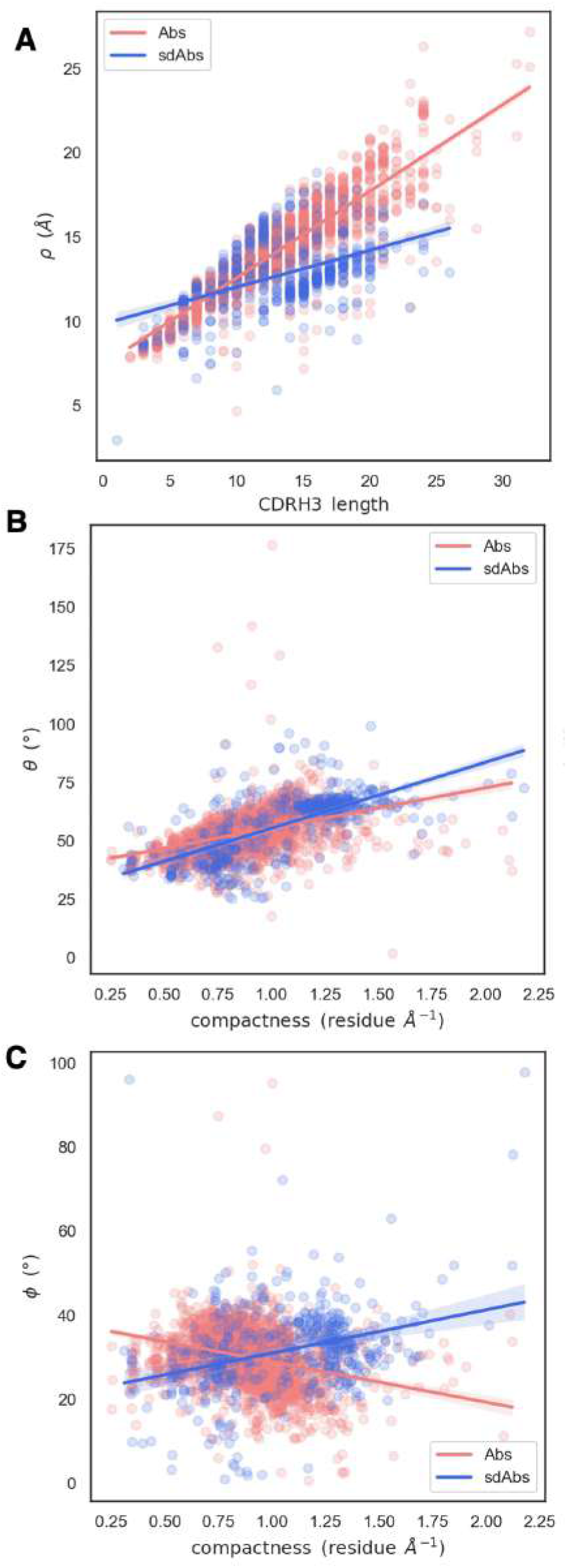
Relationships between spherical angles and compactness suggest that the paratope is stabilised by the CDR-H3 loop packing against VL domains in Abs (pink), or the rest of the VHH domain in sdAbs (blue). **(A)** Correlation between *ρ* and CDR-H3 length **(B)** Correlation between *θ* and compactness **(C)** Correlation between *ϕ* and compactness.

Compared to Ab CDR-H3 structures, sdAbs show a much wider bimodal distribution of *θ* values, with the major peak corresponding to *θ* values in excess of those observed for Ab structures, and another minor peak below the Ab distribution (Figure 7C). This indicates that the majority of sdAb CDR-H3 loops lie flat against the rest of the VHH domain, therefore folding down. We observe a slight shift in *θ* in the distribution for bound sdAbs, but note that the position of the peaks still remains stable. We conclude that sdAbs generally do not extend their CDR-H3 loops upon binding, as has previously been hypothesized. Lastly, we find near-identical values of *ϕ* for sdAbs and Abs, with sdAb *ϕ* values having a slightly wider distribution (Figure 7D).

To examine how CDR-H3 loops pack against the VH or VL domains, we analysed the relationship between the spherical angles and compactness. Both sdAb and Ab CDR-H3 loops become more compacted through an increase in *θ*: packing of the loop down towards the rest of the VH domain decreases its reach (Figure 8B). We hypothesize that this is a mechanism to stabilise the paratope structure by allowing the loop to pack against the rest of the VH domain. We also find an inverse relationship between compactness and *ϕ* for sdAbs and Abs (Figure 8C). As *ϕ* increases (as the CDR-H3 loop is horizontally oriented away from the VH domain), sdAbs show an increase in compactness whereas the opposite is true for Abs. For sdAbs, an increase in *ϕ* results in the loop extending away into empty space, whereas in Abs the loop is positioned towards the VL domain. As the presence of the VL domain provides steric hindrance, the CDR-H3 loop is forced into a conformation that orients it away from the Ab, therefore reducing compactness and increasing reach.

### SdAbs establish more interactions with their epitope per paratope residue than Abs

Our results thus far demonstrate that there are differences between the paratopes of sdAbs and Abs. But, our results also find only limited differences between the epitopes of the two modalities.

We have shown that for our datasets, Abs and sdAbs are able to bind similarly-sized epitopes, despite sdAbs paratopes being smaller. In order to investigate how this is achieved, we compare the interactions observed within binding sites. We find that, normalising for the size of the paratope, per paratope residue, sdAbs establish significantly more interactions than Abs (Figure 9). This suggests that sdAbs establish a similar binding affinity to Abs by each paratope residue having an increased number of interactions with the epitope.

**Figure 9.**
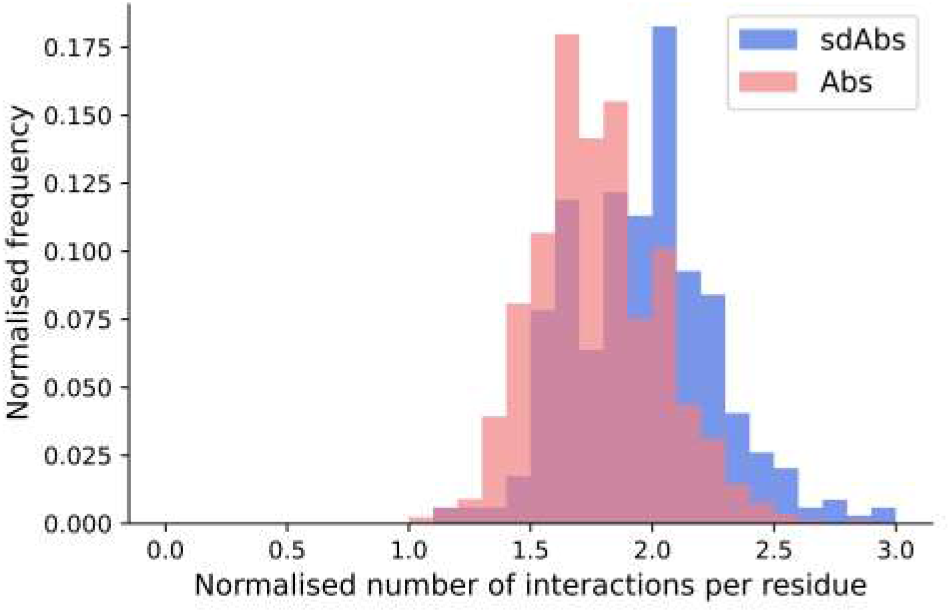
The distributions of the number of interactions initiated by sdAbs (blue) and Abs (pink) paratopes demonstrate that sdAb paratopes establish significantly more interactions per residue than Ab paratopes. Comparing the number of interactions from sdAbs to Abs, normalised for paratope size, we find a mean increase of 0.19.

### Hydrophobic interactions dominate both sdAb-Ag and Ab-Ag complexes

As well as the number of interactions, the types of interactions established between the antigen and the antibody in sdAbs and Abs were compared. All interatomic interactions between the interaction-defined epitope and paratope residues were considered. Each type of interaction was counted individually if an atom-atom complex established more than one interaction type (see Methods for full details).

In terms of interactions arising from the CDR loops, very similar types are observed (Figure 10A), whilst for the framework regions involved in binding, we see an increase in hydrophobic interactions for sdAbs compared to Abs and the VH domain of Abs alone (Figure 10B).

**Figure 10.**
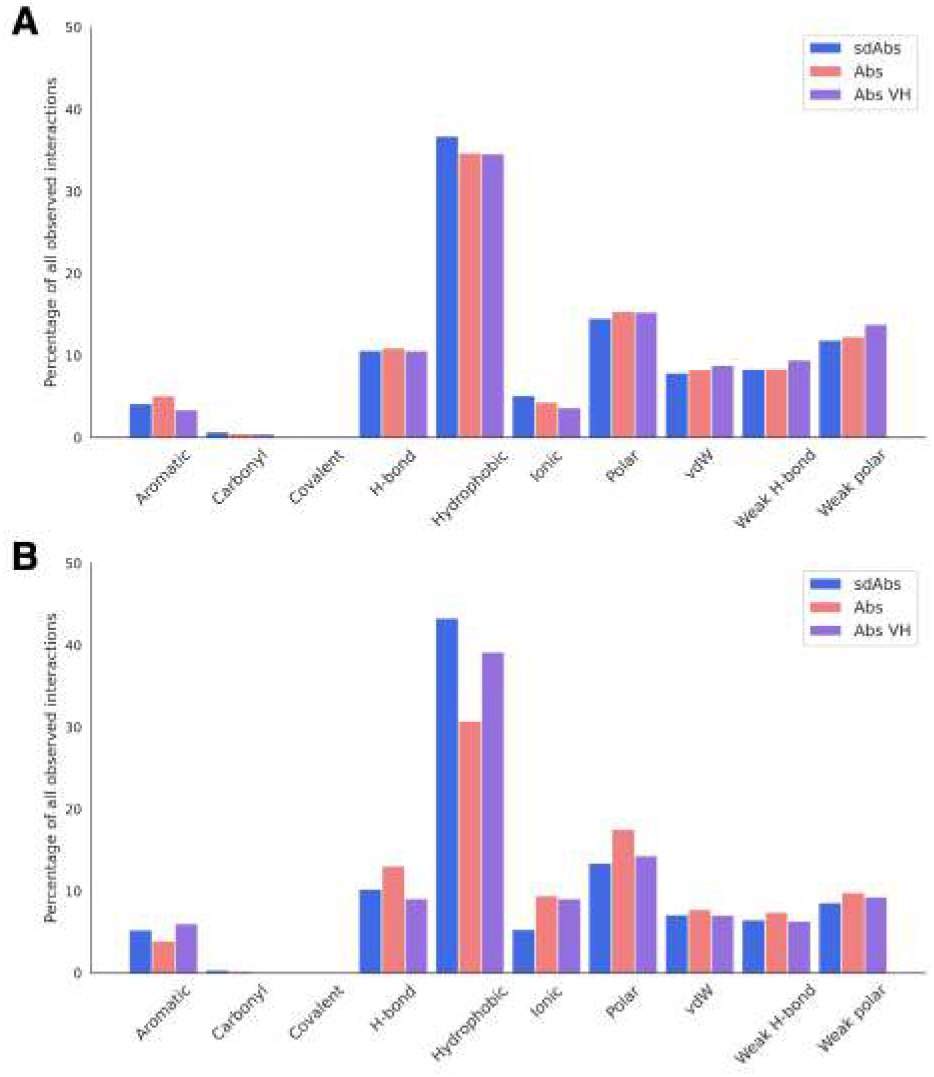
Hydrophobic interactions dominate across sdAb-Ag and Ab-Ag complexes. Total occurrences as a percentage of all interaction types observed for the **(A)** CDR loops and **(B)** the framework regions. Results for sdAbs are shown in blue, Abs are shown in pink and the VH domain of Abs are shown in purple.

### CDR-H3 and framework residues are of increased importance for interactions in the sdAb:Ag complex

In our data, we see the expected dominance of the CDR-H3 loop in binding (Supplementary Figure S16). However there are a small number of examples where the CDR-H3 loop contributes zero interactions (Figure 11). We found that there are significantly more interactions contributed from the CDR-H3 in sdAbs than Abs (Supplementary Figure S17A) even after normalising for CDR-H3 length (Supplementary Figure S17B) and that in sdAbs, there was a significantly greater contribution from the CDR-H3 residues both in terms of contributing residues to the paratope and contributing interactions (Supplementary Figure S17C, S17D). When comparing the paratope of sdAbs only to the paratope residues from the Ab VH domain, again significant differences are found (Supplementary Figure S17E, S17F). These results show that the highly variable CDR-H3 loop is even more dominant in sdAbs than in Abs. This, however, is not the only difference: we also observe that the paratopes of sdAbs tend to contain a smaller proportion of CDR residues than Abs (Figure 12, Supplementary Figure S18), from which we can infer that sdAbs show greater inclusion of framework residues in their paratopes than Abs.

**Figure 11.**
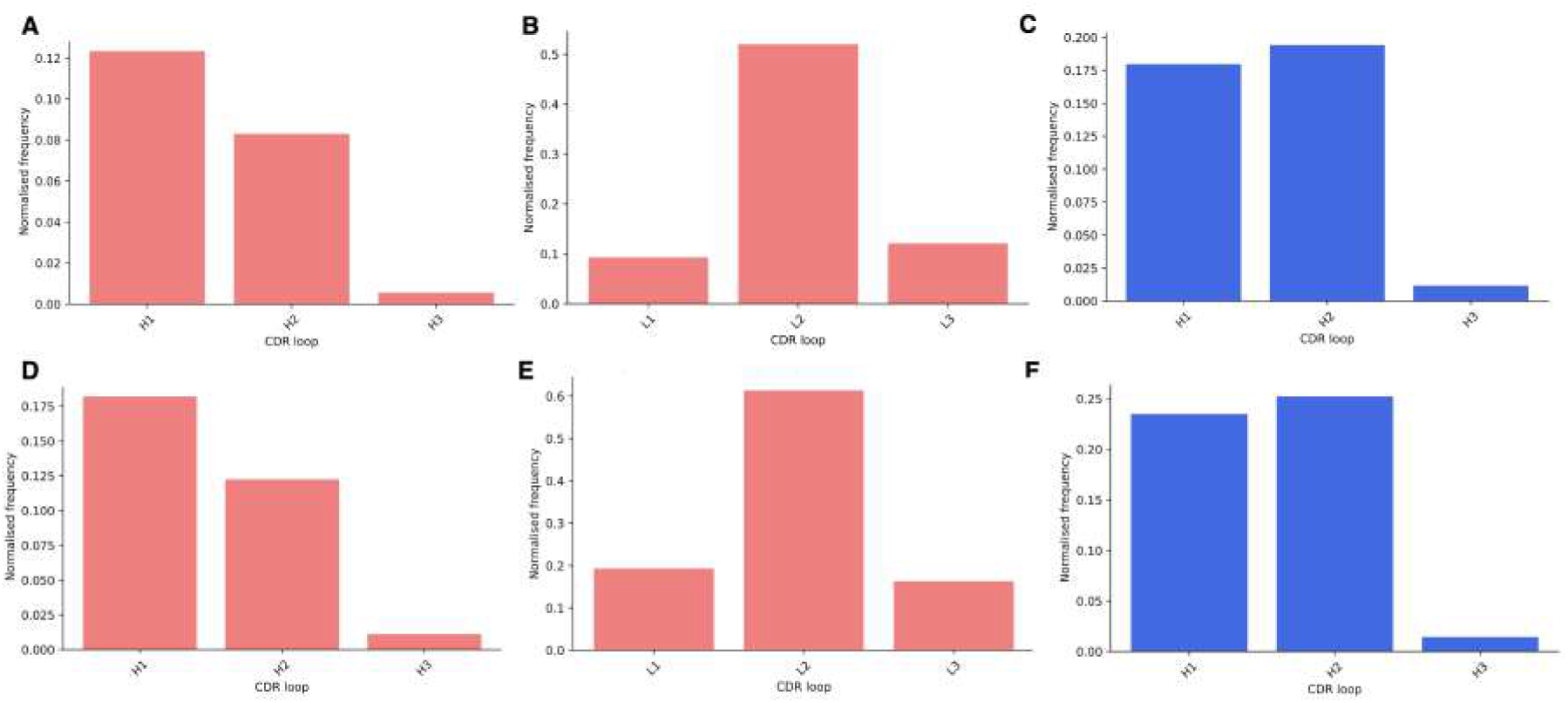
Assessing the relative contributions of each CDR loop to the paratope shows that for both sdAbs (blue) and Abs (pink), the CDR-H3 loop rarely does not contribute interactions to the paratope. Bars show the number of times a CDR loop contributes zero interactions to a paratope as a proportion of all structures in that dataset for the distance-defined (**A, B, C**) and interactions-defined (**D, E, F**) paratopes for the Abs VH, Abs VL and sdAbs respectively.

**Figure 12.**
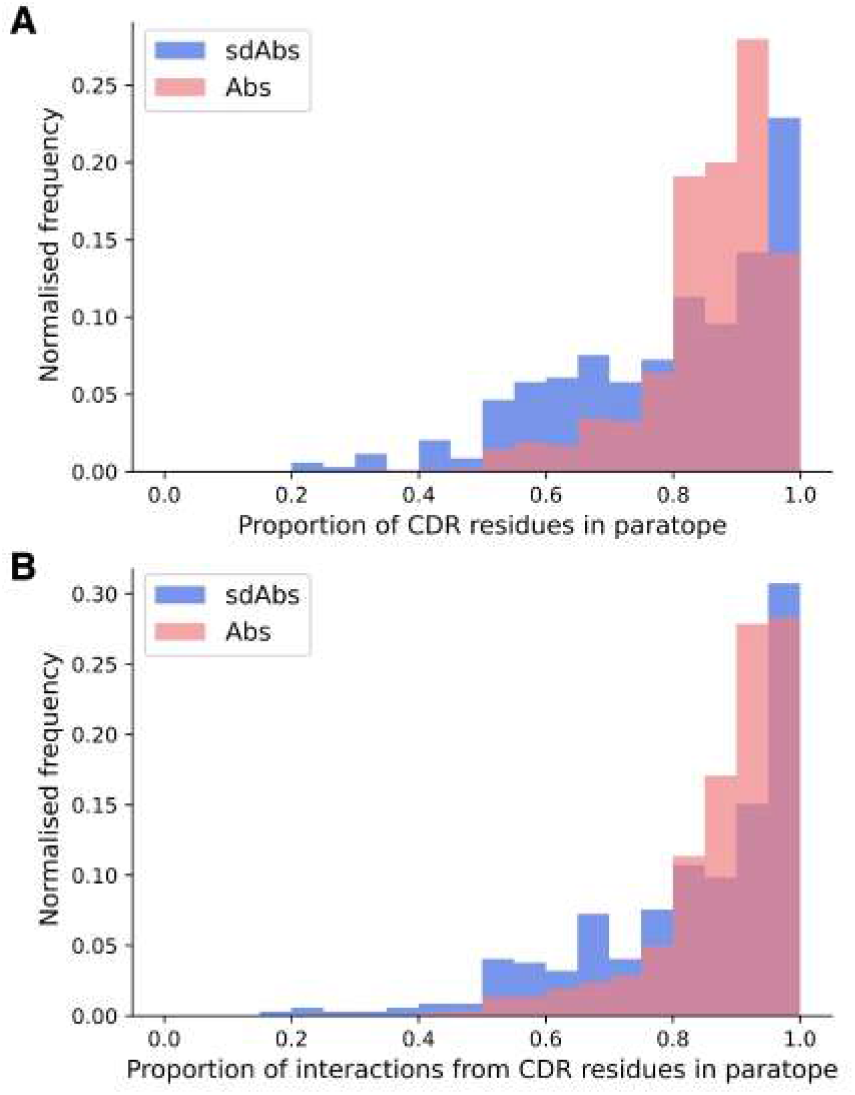
Distributions of **(A)** the proportion of CDR residues in the paratope and **(B)** the proportion of interactions from CDR residues across the whole paratope, determined per complex in the sdAbs (blue) and Abs (pink) datasets. Higher density on the lower end for the sdAb dataset (blue), compared to the Ab dataset (pink), indicates that more framework residues are involved in binding the epitope.

### Interacting framework residues are often conserved in sdAbs

Given we find that framework residues make up a larger proportion of the paratope in sdAbs than in Abs (Figure 12), we next tested if these framework residues show high variability, undergoing somatic hypermutation to improve binding, or are conserved germline residues. Framework residues observed in the interactions-defined paratope in at least 10% of the sdAb complexes were determined (Supplementary Table S3) and in descending order of frequency, include positions 66 (50.4%), 52 (31.6%), 55 (27.2%), 42 (24.1%), 50 (17.4%), 118 (15.9%), 69 (12.8%), 67 (12.8%), 40 (10.4%), and 2 (10.1%). The amino acid compositions of these identified framework positions were determined for both of the structural datasets and for the sequence datasets (Figure 13). Positions were not included if less than 5% of the structures or sequences had a residue at that position. We compare the positions found in the interactions-defined paratopes from the structural datasets to a background composition taken from the sequence datasets. The sequence logo plots (Figure 13), show similarities between the paratope composition and background particularly for positions 2, 50, 67, 69 and 118 in sdAbs. The low level of variation at these positions in sdAbs indicates they are conserved and suggests that they may not contribute to binding specificity.

**Figure 13.**
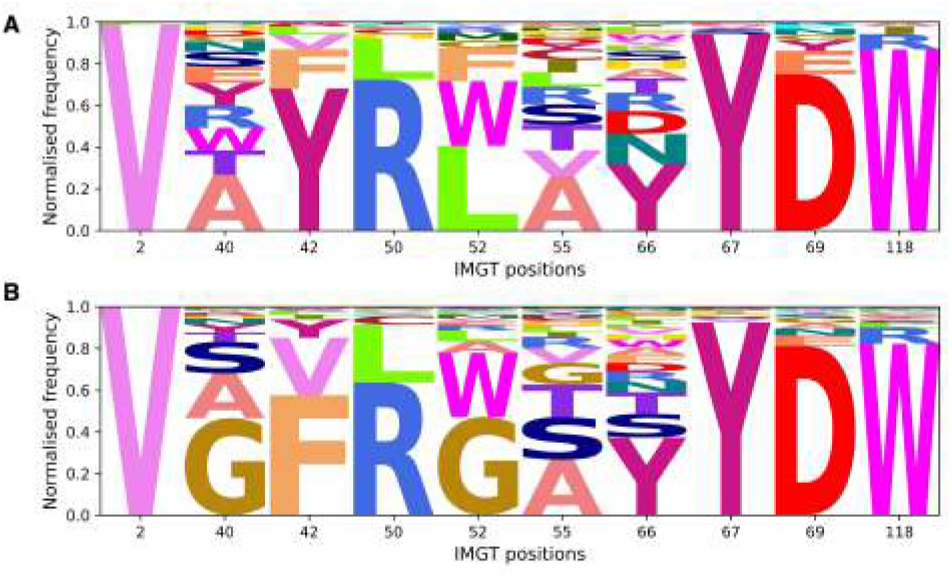
Sequence logo plots for framework positions often involved in the paratopes of Abs and sdAbs suggest that framework residues identified to often occur in the paratope are highly conserved in sdAbs. **(A)** Amino acid compositions at positions found in at least 10% of sdAbs paratopes in our sdAbs structural dataset. **(B)** Background amino acid compositions in our sdAbs sequence dataset for positions found in at least 10% of sdAbs paratopes. Positions were not included if less than 5% of sequences had a residue at the given position.

### Abs and sdAbs can bind the same epitopes but interact with them differently

Our results suggest that Abs and sdAbs can engage similar types of epitopes but use different mechanisms to do so. Here, we compare the features of an Ab (PDB ID: 6YLA) and a sdAb (PDB ID: 6WAQ) that both bind to the receptor-binding domain (RBD) of the SARS-CoV-2 spike protein, using interactions-defined binding sites. The sdAb has a longer CDR-H3 (18 residues) than the Ab (12 residues) and the sdAb paratope is smaller than that of the Ab (15 compared to 26 residues). The sdAb paratope includes framework positions 66 and 69, both of which we found to be commonly part of sdAb paratopes. The Ab paratope includes framework positions 1 from the heavy chain and position 68 from the light chain.

Despite the differences in the sdAb and Ab paratopes, they are binding a very similar epitope (Figure 14). The epitopes on the RBD that these structures bind are of a similar size (15 residues for the Ab epitope and 18 residues for the sdAb epitope).

**Figure 14.**
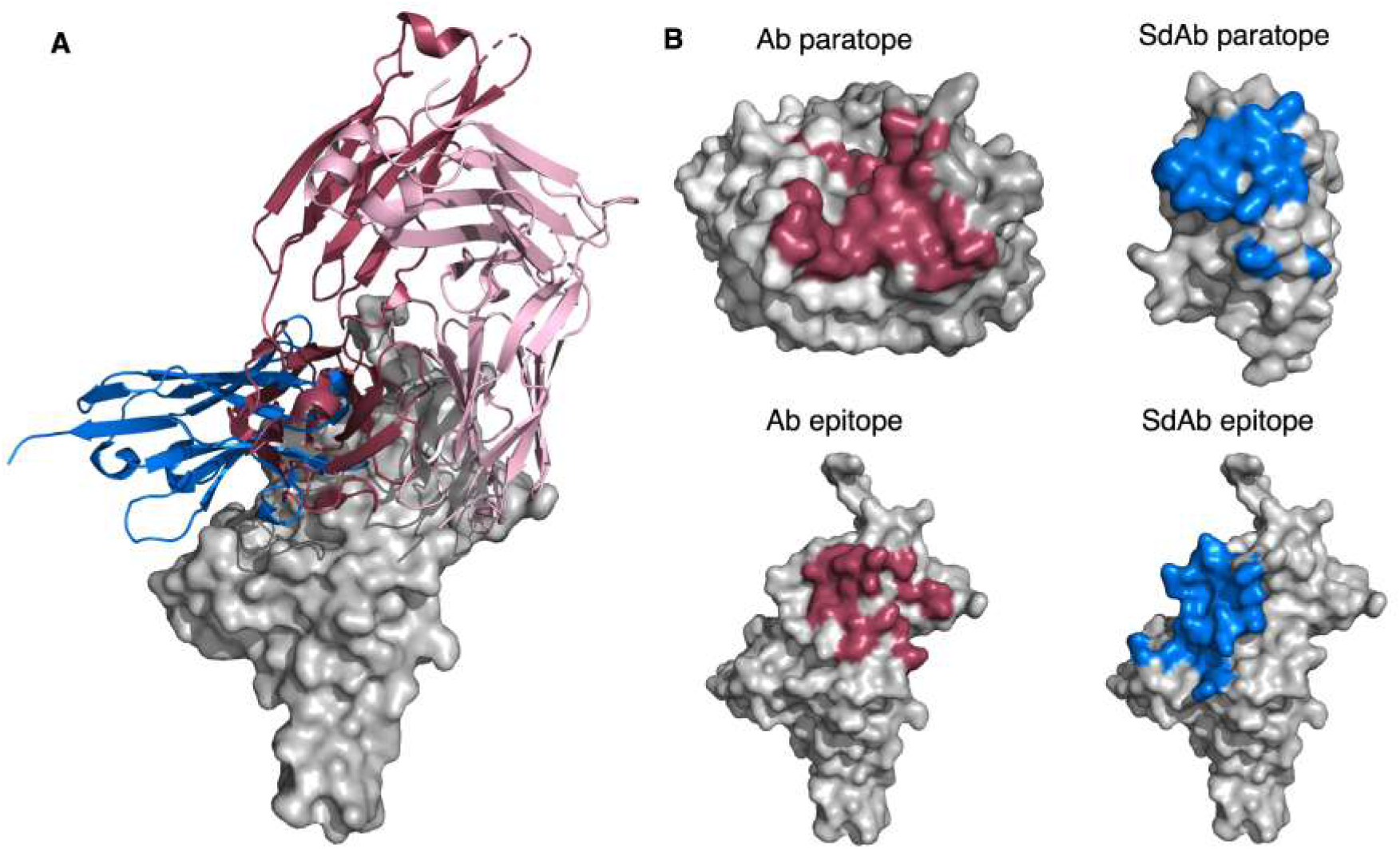
**(A)** A sdAb (PDB ID: 6WAQ) and Ab (PDB ID: 6YLA are able to bind the SARS-CoV-2 RBD with overlapping epitopes. Dark pink cartoon = Ab heavy chain, light pink cartoon = Ab light chain, blue cartoon = sdAb, grey = surface representation of the SARS-CoV-2 RBD. **(B)** Abs in general have larger paratopes than sdAbs, but sdAbs are able to bind similarly-sized epitopes as exemplified by structures 6YLA (Ab) and 6WAQ (sdAb). The surface of the Ab heavy chain is shown in dark grey and the light chain in light grey, where the blue region represents paratope residues. The surface of the sdAb is shown in light grey with the pink region representing the sdAb paratope residues. The surface of the SARS-CoV-2 antigen is shown in light grey for both the sdAb and Ab, where the Ab epitope is coloured pink and the sdAb epitope in blue. The antigen structures from each PDB were merged to create a complete image of the antigen for the sdAb.

Thirty-one total interactions occur between the Ab epitope and paratope, whilst there are twenty-nine for the sdAb binding site, however when we consider the size of the paratope, this results in an average of 1.9 interactions per paratope residue for the sdAb, compared to 1.2 per Ab paratope residue. In addition, the CDR-H3 has increased importance for the sdAb binding activity. For the Ab, 6 out of the 26 residues in the paratope come from the CDR-H3 loop, whereas for the sdAb, it is 9 out of 15.

## Discussion

In this study, we compared the binding sites of sdAbs and Abs to assess whether these two modalities may be suited to different types of epitopes. Overall we find that the paratopes of sdAbs and Abs show similarities in their amino acid compositions, but otherwise have distinguishable characteristics. Paratopes of sdAbs tend to be smaller, the CDR conformations observed are different between sdAbs and Abs, and sdAbs tend to have longer CDR-H3 loops than their Ab counterparts. These results are all consistent with previous studies on smaller datasets (*9, 12, 18*).

These differences in their paratopes led to the expectation that Abs and sdAbs would bind distinct types of epitopes. However, we find that, apart from the epitopes of Abs being slightly more linear than those of sdAbs, the epitopes targeted by sdAbs and Abs cannot be easily distinguished. SdAbs and Abs target epitopes of similar size, similar amino acid compositions and similar shape. There are several suggestions in the literature that the longer CDR-H3 loop of a sdAb means it can interact with more concave epitopes by protruding into the cavity (*13*–*16*). Henry and MacKenzie (2018) (*17*) stress that despite individual case studies supporting this hypothesis, the evidence that sdAbs prefer concave epitopes is limited and it is unknown whether this is a general trend across sdAbs. We find that overall, for our datasets, the epitopes targeted by sdAbs are not more concave than epitopes targeted by Abs. Furthermore, we find no correlation between CDR-H3 loop length and epitope concavity. It should be noted that our findings are limited to represent the epitopes of solved structures only, and it is possible that they do not translate to the wider epitope diversity recognised by Abs or sdAbs in nature.

However, these results are supported by our finding that Ab and sdAb CDR-H3 loops show differences in their orientation relative to the rest of the supporting VH/VL or VHH domain. We find that sdAb CDR-H3 loops are more compacted than Ab loops and are often found packed against the rest of the VHH domain. For Abs, orientation of the CDR-H3 away from the VH domain leads to its positioning towards the VL domain. As the presence of the VL domain provides steric hindrance, the CDR-H3 loop is forced into a conformation that orients it away from the Ab, therefore reducing compactness and increasing reach. In contrast, for sdAbs, orientation of the CDR-H3 away from the VH domain leads to positioning towards empty space and therefore packing against the rest of the VHH domain. These results offer a possible explanation for our observation that the longer CDR-H3 loops of sdAbs do not necessarily target deeper epitopes.

In addition, we observe that framework residues are more often observed in the paratopes of sdAbs. The importance of framework residues in sdAbs has been indicated in several studies (*9, 12, 42, 43*). This increase in framework residues is likely related to their increased accessibility due to the lack of the VL domain. Indeed, our results show that most of the framework positions observed in more than 10% of the sdAbs paratopes are frequently observed in the VH-VL interface of Abs (*44*). Most of the framework positions commonly involved in binding in sdAbs belong to FR2, which is identified by both Zavrtanik *et al*. (2018) (*9*) and Mitchell and Colwell (2018a) (*12*) as an important region for antigen binding. The majority of our identified potential paratope framework residues appear to be highly conserved. Our findings that sdAb CDR-H3 loops often pack against the VHH domain, and that FR2 residues are often conserved, is in agreement with that of Sang *et al*. (2022) (*43*), who find that the longer CDR-H3 loops of sdAbs can fold back to interact with FR2 residues.

Finally, we also find that despite tending to have smaller paratopes, sdAbs are able to target similarly-sized epitopes to Abs. This may be explained by our finding that the CDR-H3 loops of sdAbs make a significantly greater number of interactions with the epitope per loop residue than those of Abs, even after normalising by loop length. Given that these may include conserved framework residues, that will contribute to binding affinity but not specificity, this raises important questions over the specificity of the sdAb binding site, as well as having implications for engineering therapeutics.

## Conclusions

Overall, this study highlights structural characteristics of sdAbs pertinent to the design and engineering of sdAb therapeutics, and calls attention to the need for additional criteria when deciding on the best modality for a particular epitope.

## Acknowledgements

This work was supported by the Engineering and Physical Sciences Research Council (grant number EP/S024093/1), Twist Bioscience and the Wellcome Trust (grant number 102164/Z/13/Z). The authors thank Claire Marks and Katya Putintseva from LabGenius for their work on the project.

