## Supplementary Material for "A comparison of the binding sites of antibodies and single-domain antibodies"

### Supplementary Text

##### Text S1

After creating the structural datasets as described in the Methods section, four sdAb complexes and seven Ab complexes were removed. The removed sdAb PDB entries are 7D30, 6UL6, 4HIJ and 5OMM. The first two consist both of two individual sdAb molecules identified by the same chain identifier, resulting in an uncommonly large number of interactions between the antigen and the antibody. PDB entry 4HIJ was removed because it was erroneously annotated as a sdAb. PDB entry 5OMM is lacking complete CDRs. These latter two entries were removed after studying complexes for which the fraction of CDR residues in the paratope was remarkably low. For the Ab dataset, the PDB entries 2H32, 4NZR, 4NZT, and 1XIW, and the complexes including antibody chains KJ and ML in PDB entry 5MHR were removed. The first three entries were highlighted because the number of interactions was remarkably high and the fraction of the CDR residues in the paratope remarkably low. PDB entry 2H32 does not include a complete full-length antibody. PDB entries 4NZR and 4NZT were removed because both include a protein binding to an antibody, instead of an antibody binding to an antigen. PDB entry 1XIW contains two Abs which are both sdAbs and are thus incorrectly annotated in the SAbDab database: there was a similar case for complexes in PDB entry 5MHR . The fraction of the number of interactions in the paratope that were established by CDR residues was low for the complexes of PDB entry 1XIW. The fraction of CDR residues in the paratope was low for the complexes of PDB entry 5MHR.
Arpeggio failed to run for two PDB entries, even after cleaning the PDB files. These PDB entries are 4XWO and 5K9Q and contained antibodies. The Ab dataset containing the Arpeggio-defined epitopes and paratopes therefore has a size of 890 complexes.

**Text S2**

We analysed the general orientation of CDR-H3 loops of Abs and sdAbs by examining their centers of geometry in reference to an $\mathbb{R}^3$ coordinate system. We describe the center of geometry of the CDR-H3 loop using spherical coordinates. To orient the coordinate system identically in all examined structures, we used the following fitting procedure.

Let $\mathbf{c}$ be the centre of geometry of the backbone atoms of IMGT positions 102, 103, 118, and 119. Let $\mathbf{p}$ be the first principal component vector of the anchor positions which minimises the distance to the C$\alpha$ atoms of IMGT position 103 when extended from $\mathbf{c}$. Let $\mathbf{a}$ be the centre of geometry of all VH anchor positions (we take the anchors to be the three residues on either side of the loop as defined by the IMGT numbering scheme.). Let $\mathbf{\Pi}$ be the plane that crosses $\mathbf{c}$ whose normal vector is $\mathbf{p}$. Therefore for all $\mathbf{r} = \langle x, y, z \rangle$,

$$\mathbf{p} \cdot (\mathbf{r} - \mathbf{c}) = 0 \tag{S1}$$

$$\mathbf{p} \cdot \mathbf{r} = \mathbf{p} \cdot \mathbf{c}\,. \tag{S2}$$

We calculate the shortest distance of any point $\mathbf{r}$ from $\mathbf{\Pi}$ using

$$d(\mathbf{r}) = \frac{\mathbf{p} \cdot \mathbf{r} - \mathbf{p} \cdot \mathbf{c}}{||\mathbf{p}||}\,. \tag{S3}$$

Let $\mathbf{u} = \mathbf{a_\Pi} - \mathbf{c}$, where $\mathbf{a_\Pi}$ is the orthogonal projection of $\mathbf{a}$ onto $\mathbf{\Pi}$, given by

$$\mathbf{a_\Pi} = \mathbf{a} - d(\mathbf{a}) \frac{\mathbf{p}}{||\mathbf{p}||}\,. \tag{S4}$$

Let $\mathbf{v}$ be a vector orthogonal to both $\mathbf{u}$ and $\mathbf{p}$, such that $\mathbf{v}$ the resultant coordinate system is left-handed,

$$\mathbf{v} = \underset{\mathbf{x} \in \{\mathbf{u} \times \mathbf{p}, -\mathbf{u} \times \mathbf{p}\}}{\operatorname{argmax}} \begin{vmatrix} x^{(1)} & u^{(1)} & p^{(1)} \\ x^{(2)} & u^{(2)} & p^{(2)} \\ x^{(3)} & u^{(3)} & p^{(3)} \end{vmatrix}\,. \tag{S5}$$

Given the vectors $\mathbf{u}$, $\mathbf{v}$, and $\mathbf{p}$, such that $\mathbf{u} \perp \mathbf{v} \perp \mathbf{p}$, we aim to describe the CDR-H3 backbone atomic positions using spherical coordinates.

Let $\mathbf{h}$ be the centre of geometry of the CDR-H3 backbone positions, as defined by the IMGT numbering scheme. To describe $\mathbf{h}$ relative to $\mathbf{c}$ using spherical coordinates, $\mathbf{h_c}(\rho, \theta, \phi)$, we define $\rho = ||\mathbf{h} - \mathbf{c}||$. $\theta$ describes the elevation or polar angle, and $\phi$ describes the azimuth. $\theta$ decreases as the CDR-H3 loop projects directly up and away from the VH. $\phi$ increases as the CDR-H3 loop points out from the VH. In the case of antibodies, this refers to a CDR-H3 loop that is oriented more towards the VL domain.

We use $\mathbf{p}$ as the polar reference vector. Thus,

$$\theta = \cos^{-1} \frac{(\mathbf{h} - \mathbf{c}) \cdot \mathbf{p}}{||\mathbf{h} - \mathbf{c}||\; ||\mathbf{p}||}\,. \tag{S6}$$

Lastly, we use $\mathbf{u}$ as the azimuthal reference vector. Therefore, $\phi$ will be the angle between $\mathbf{u}$ and $\mathbf{h_\Pi} - \mathbf{c}$, where $\mathbf{h_\Pi}$ is the orthogonal projection of $\mathbf{h}$ onto $\mathbf{\Pi}$,

$$\mathbf{h_\Pi} = \mathbf{h} - d(\mathbf{h}) \frac{\mathbf{p}}{||\mathbf{p}||} \tag{S7}$$

and

$$\phi = \cos^{-1} \frac{(\mathbf{h_\Pi} - \mathbf{c}) \cdot \mathbf{u}}{||\mathbf{h_\Pi} - \mathbf{c}||\; ||\mathbf{u}||}\,. \tag{S8}$$

In order to consistently measure the azimuth angle, we calculate

$$\begin{vmatrix} h_\Pi^{(1)} & u^{(1)} & p^{(1)} \\ h_\Pi^{(2)} & u^{(2)} & p^{(2)} \\ h_\Pi^{(3)} & u^{(3)} & p^{(3)} \end{vmatrix}\,.$$

As we aim to measure the clockwise displacement angle using $\mathbf{p}$ as a reference, we use the calculated $\phi$ as the azimuth angle when the determinant is positive. Else we define the azimuth reference angle as $2\pi - \phi$.

#### Supplementary Tables

**Table S1.** The number of complexes (NoC) remaining in the sdAb structural dataset and the Ab structural dataset are shown after performing each filtering step. PDB entries containing antibodies in complex with antigens were selected. All Ab-Ag complexes having at least one CDR residue as binding residue were identified. Next, outliers were removed. Only structures containing the antibody in complex with the protein antigen were selected, using the annotations provided by SAbDab. Moreover, complexes with an epitope and/or paratope size smaller than seven residues were removed. Clustering with cd-hit was performed using a 95% cut-off to filter on CDR sequence identity. Complexes were reintroduced if their epitope similarity with any other selected complex in the same resulting cd-hit cluster was less than 75%.

| **Filtering steps** | **Total sdAbs** | **Total Abs** |
|---|---|---|
| Selected PDB entries | 411 | 1260 |
| NoC close CDR residue | 1044 | 1751 |
| NoC removed outliers | 1038 | 1045 |
| NoC filtered SAbDab | 816 | 1703 |
| NoC filtered binding site size | 781 | 1699 |
| NoC filtered CDR sequence identity | 309 | 792 |
| NoC reintroduced non-identical epitopes | 345 | 892 |

**Table S2.** Greater species variation is observed in Abs. The species assigned to the antibodies in both the sdAb structural dataset and the Ab structural dataset. Species are derived from the VH domain of the antibodies as stored in the SAbDab database. Note, these annotations are not always correct, especially for the sdAbs due to the fine line between humanised VHHs and camelidised VHs.

| **Species** | **sdAbs** | **Abs** |
|---|---|---|
| *Lama glama* | 152 | 4 |
| *Vicugna pacos* | 94 | |
| *Homo sapiens* | 19 | 557 |
| *Camelus dromedarius* | 26 | |
| *Camelidae* | 10 | |
| None | 9 | 15 |
| Synthetic construct | 16 | 6 |
| *Mus musculus* | 1 | 250 |
| *Lama pacos* | 1 | |
| *Camelidae* mixed library | 5 | |
| *Lama* | 5 | |
| *Camelus bactrianus* | 5 | |
| *Norovirus* | 1 | |
| Unidentified | 1 | 2 |
| *Rattus norbegicus* | | 14 |
| *Oryctolagus cuniculus* | | 11 |
| *Pan troglodytes* | | 1 |
| Synthetic | | 1 |
| *Gallus gallus* | | 2 |
| *Mus* | | 2 |
| *Mus musculus, Homo sapiens* | | 8 |
| Influenza A virus | | 2 |
| *Homo sapiens*, synthetic construct | | 2 |
| *Rattus* | | 2 |
| *Macaca mulatta* | | 7 |
| *Cricetulus migratorius* | | 3 |
| *Rattus rattus* | | 2 |
| Other sequences | | 1 |

**Table S3.** The most important framework residues in sdAbs for antigen-binding observed in FR2. Framework IMGT positions, and their corresponding framework region, that are observed in the paratope for at least 10% of sdAbs are shown. The table is sorted on the observed percentage in sdAbs. The observed percentage in the Ab indicates that only positions 66 and 55 are observed in the paratope for more than 10% of the Abs. Positions in bold tend to lie in the VL-VH interface according to a study by Raybould *et al.* (2021)(*44*).

| **Position** | **Region** | **Observed percentage sdAbs** | **Observed percentage Abs** |
|---|---|---|---|
| 66 | FR3 | 50.4 | 49.0 |
| 52 | FR2 | 32.6 | 6.00 |
| 55 | FR2 | 27.2 | 34.0 |
| 42 | FR2 | 24.1 | 0.00 |
| 50 | FR2 | 17.4 | 0.00 |
| 118 | FR4 | 15.9 | 0.10 |
| 69 | FR3 | 12.8 | 5.50 |
| 67 | FR3 | 12.8 | 3.40 |
| 40 | FR2 | 10.4 | 4.80 |
| 2 | FR1 | 10.1 | 4.80 |

### Supplementary Figures

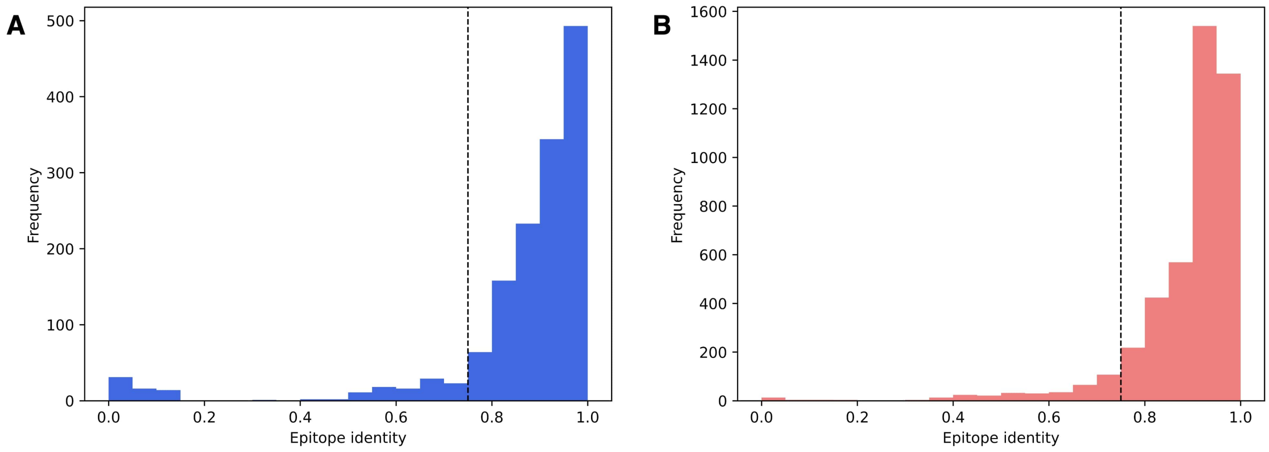


**Figure S1.** An epitope identity cut-off of 75% (dotted line) was used for reintroducing complexes in the dataset. The epitope identity of all the complexes within each cluster determined by cd-hit is shown for **(A)** sdAbs and **(B)** Abs.

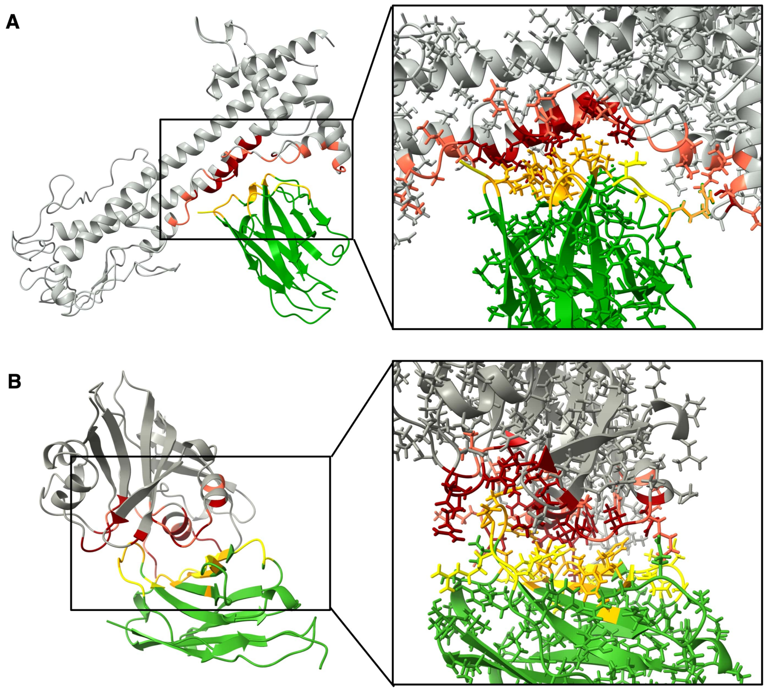


**Figure S2.** Stricter binding site definitions are obtained by using the program Arpeggio. **(A)** Complex between a sdAb (chain C) and an antigen (chain A) corresponding to PDB entry 7AQX. **(B)** Complex between a sdAb (chain B) and an antigen (chain A) corresponding to PDB entry 6QUP. The antigen is coloured grey, the positions exclusively annotated by distance-defined epitope is pink, and positions in both distance-defined epitope and Arpeggio-defined epitope dark red. The antibody is coloured green, the positions exclusively annotated by distance-defined paratope is yellow, and positions in both distance-defined paratope and Arpeggio-defined paratope orange. The pink- and yellow-coloured residues are the contacts. The dark red- and orange-coloured residues are identified as real interactions.

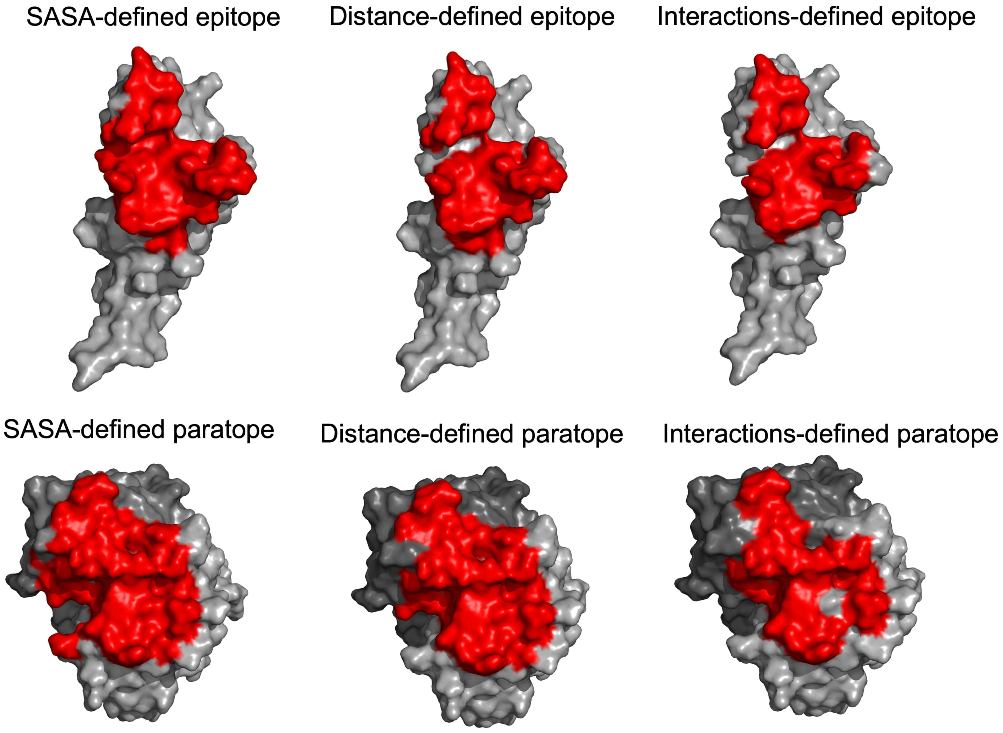


**Figure S3.** Visualising the differences between the SASA-defined, distance-defined and interactions-defined epitopes and paratopes. PDB ID: 6AZZ. For the epitope representations above, the epitope surface is red, whilst the antigen is grey. For the paratope representations below, the paratope surface is red, whilst the heavy chain of the antibody is dark grey and the light chain is light grey.

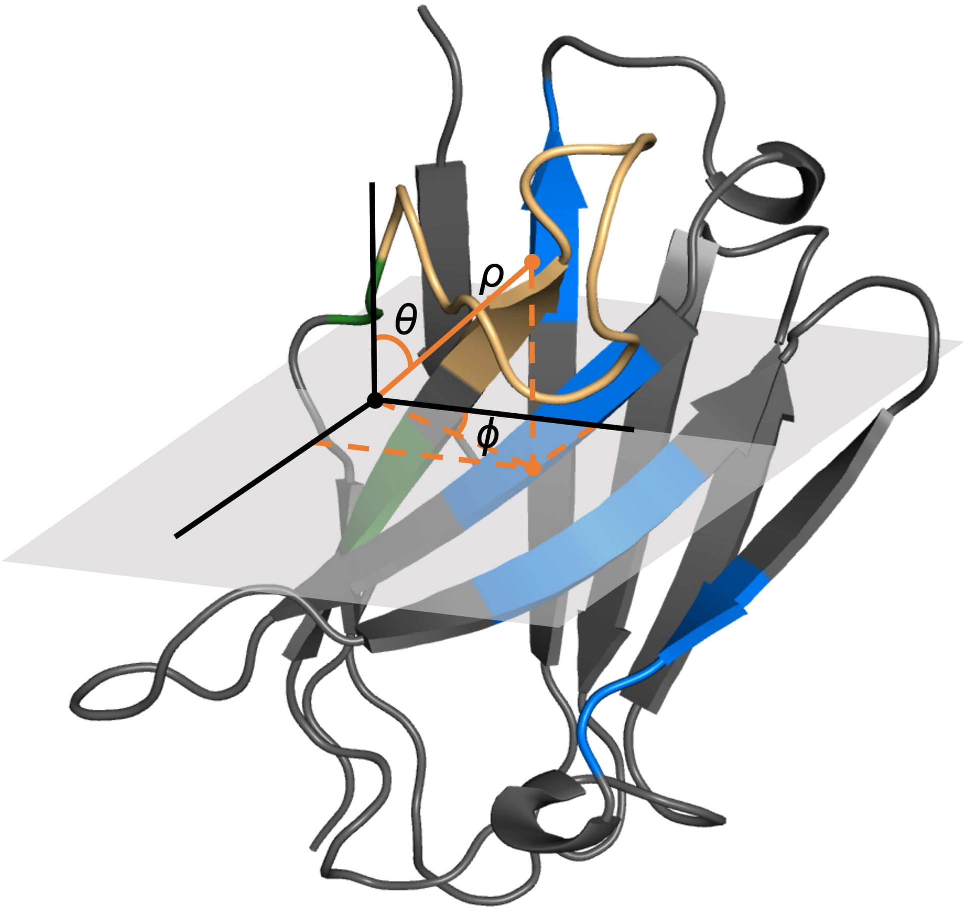


**Figure S4.** A visualisation of the coordinate system used to determine the orientation of the CDR-H3 loops, where $\rho$ describes the reach of the CDR-H3 loop away from the rest of the VH domain, $\phi$ gives an indication of whether the CDR-H3 loop is horizontally oriented towards the rest of the VH domain or away from it, and $\theta$ gives a measure of the elevation of the loop.

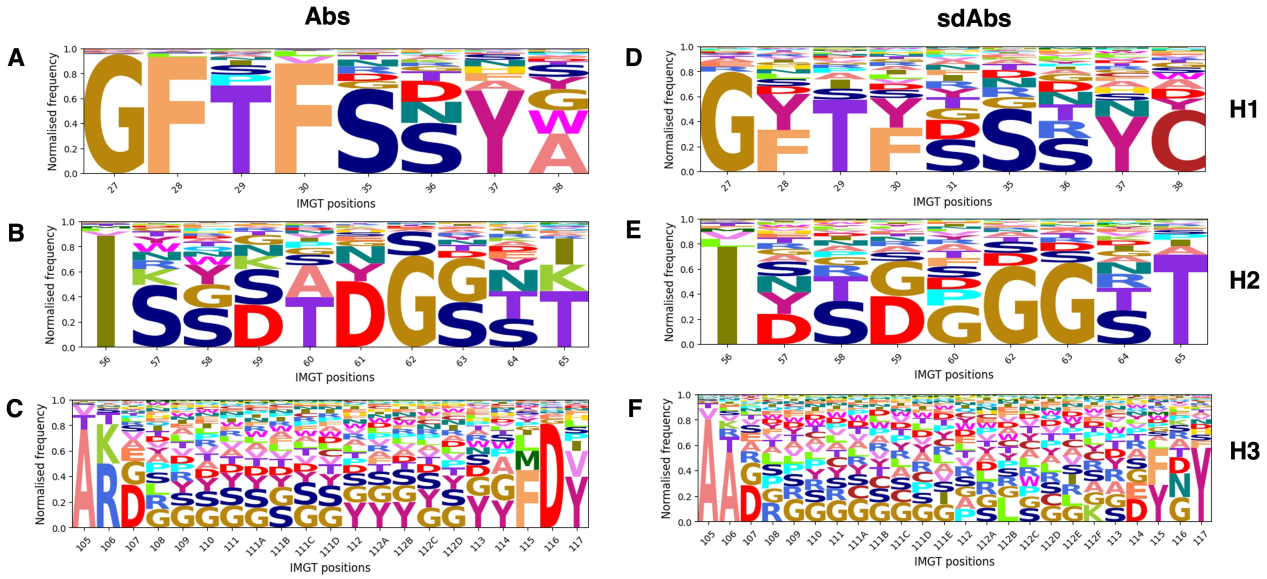


**Figure S5.** Sequence logo plots of the amino acid composition at each position of each of the CDR loops CDR-H1 (**A, D**), CDR-H2 (**B, E**) and CDR-H3 (**C, F**) shows similarities for sdAbs sequences (all belonging to germline IGHV3) and Abs IGHV3 sequences. Sequence logo plots were generated by determining the proportions of each amino acid in each position across all sequences. All positions that occurred in less than 5% of sequences have been removed.

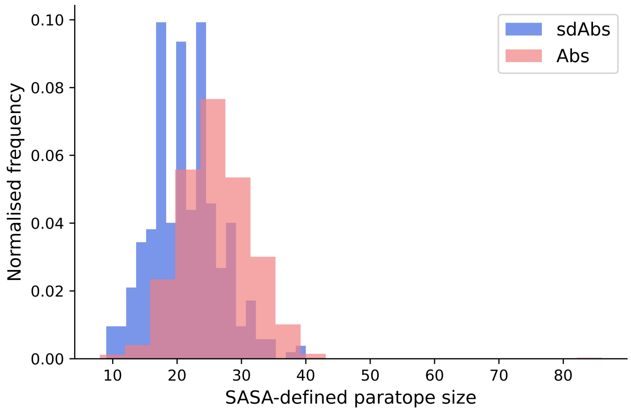


**Figure S6.** Comparing the distributions of paratope size for sdAbs and Abs complexes using the SASA-defined paratope shows that the sdAbs paratopes are significantly smaller by 4.9 residues on average (p-value « 0.01)

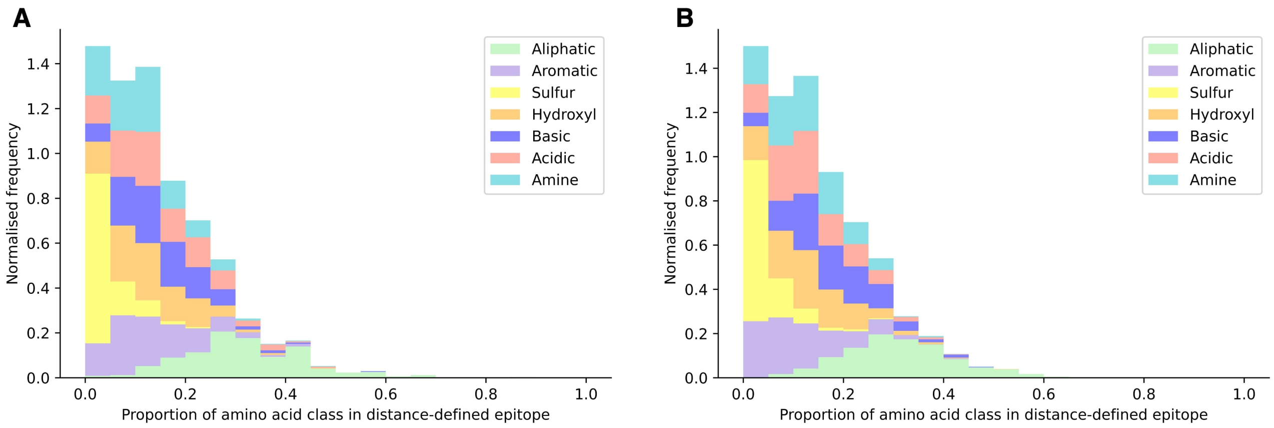


**Figure S7.** Distance-defined epitopes of sdAbs **(A)** and Abs **(B)** show similar amino acid compositions. The seven classes are: aliphatic, aromatic, sulfur, hydroxyl, basic, acidic and amine. The p-values obtained by the two-sided permutation t-test after bootstrap re-sampling on every class respectively are: 0.63, 0.015, 0.64, 0.76, 0.00060, 0.080, 0.25.

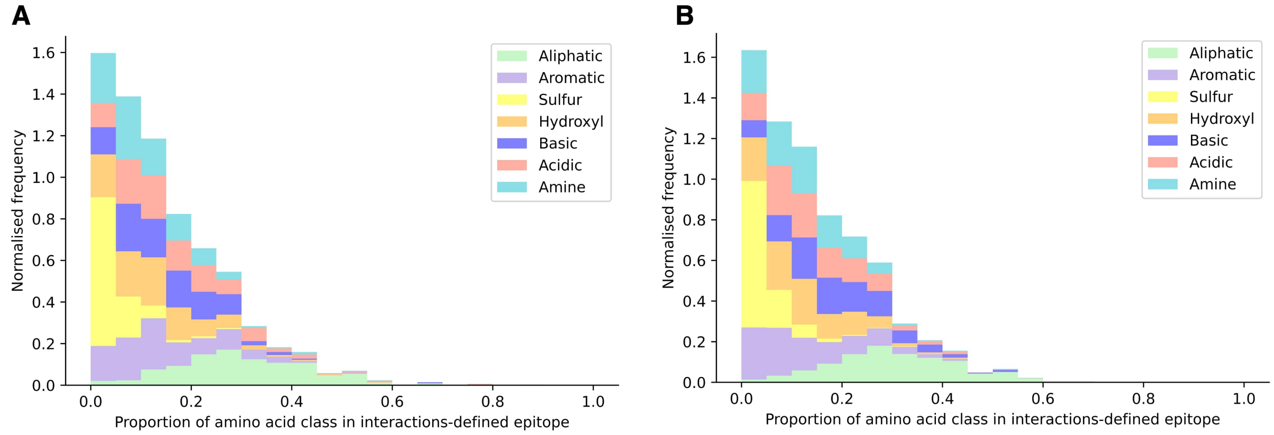


**Figure S8.** Interaction-defined epitopes of sdAbs **(A)** and Abs **(B)** show similar amino acid class compositions. The seven classes are: aliphatic, aromatic, sulfur, hydroxyl, basic, acidic and amine. The p-values obtained by the two-sided permutation t-test after bootstrap re-sampling on every class are: 0.11, 0.027 , 0.93, 0.31, 0.0014, 0.058, 0.74.

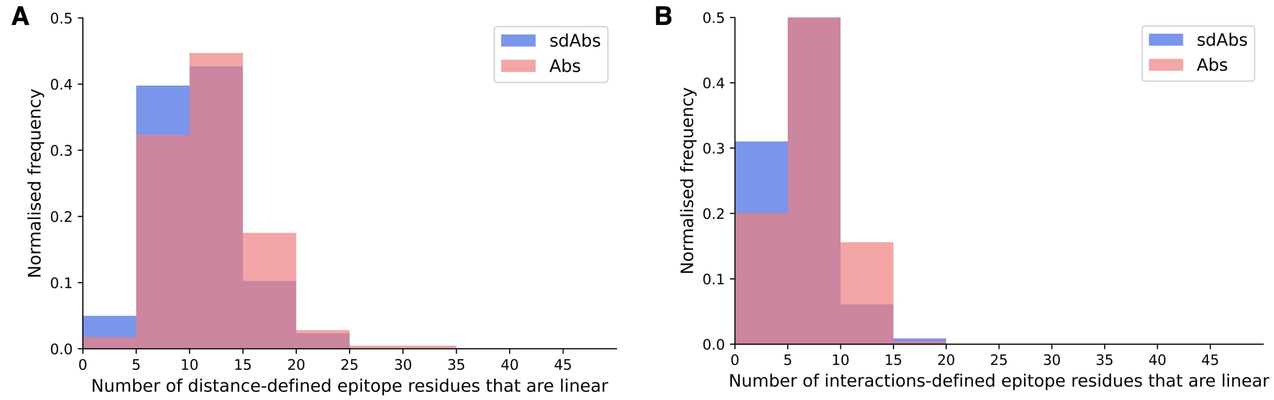


**Figure S9.** The epitopes of Abs are relatively more linear than the epitopes of sdAbs. Distributions of raw counts of linear residues for epitopes of Abs (pink) and sdAbs (blue) for the **(A)** distance-defined epitopes and **(B)** interactions-defined epitopes.

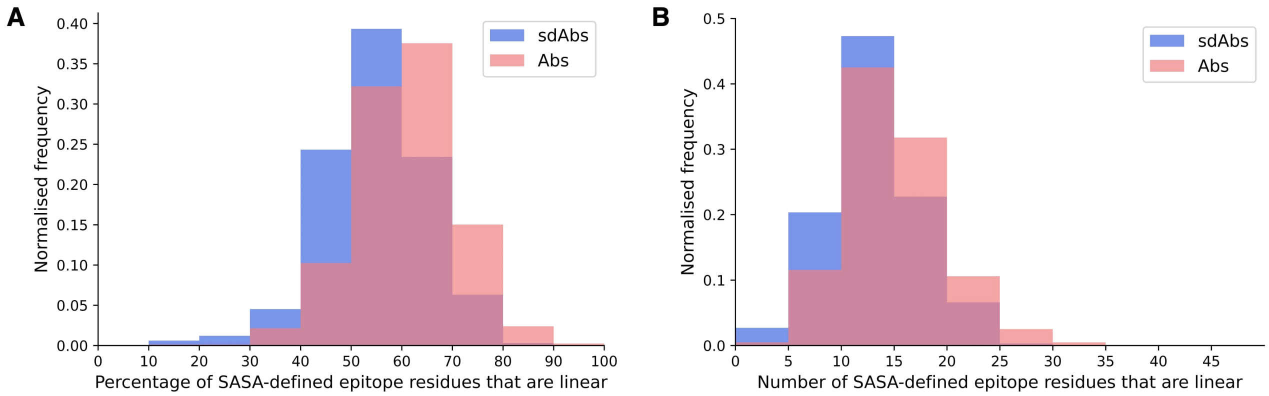


**Figure S10.** The SASA-defined epitopes of Abs are relatively more linear than the SASA-defined epitopes of sdAbs. **(A)** Distributions of the percentage of linear residues for epitopes of Abs (pink) and sdAbs (blue). The unpaired mean difference between sdAbs and Abs was 6.3% (p-value « 0.01). **(B)** Distributions of raw counts of linear residues for epitopes of Abs and sdAbs. The unpaired mean difference between sdAbs and Abs was 1.6 residues (p-value « 0.01).

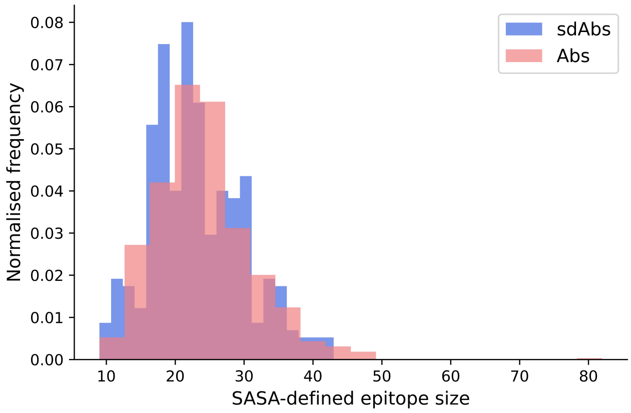


**Figure S11.** Comparing the distributions of epitope size for sdAbs and Abs complexes using the SASA-defined paratope shows that there is no significant difference in epitope size for sdAbs and Abs.

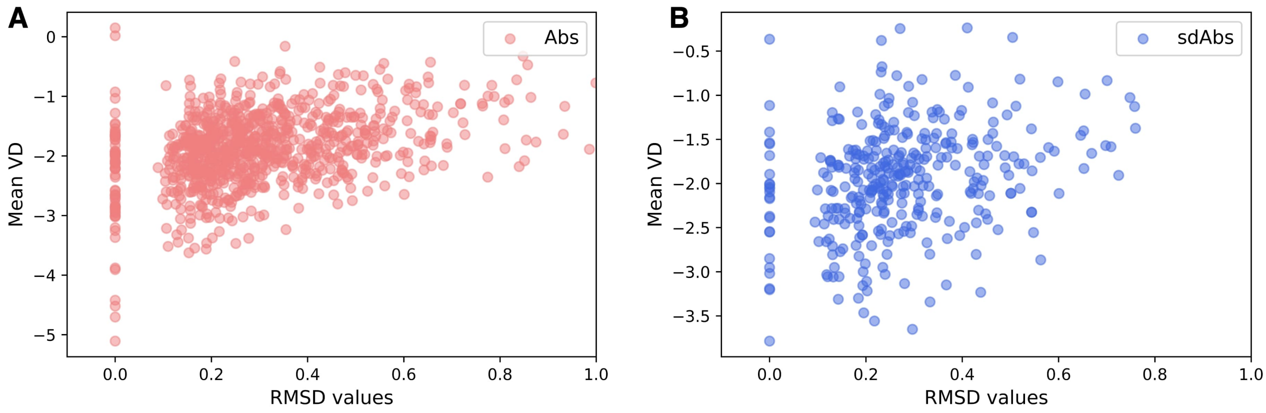


**Figure S12.** Validating the alpha shape representations of our structural datasets. No correlation of curvature of epitope surface against RMSD of mesh vertices coordinates compared to PDB coordinates shows that alpha parameter has been set such that the alpha shape accurately represents the geometry of the protein surface.

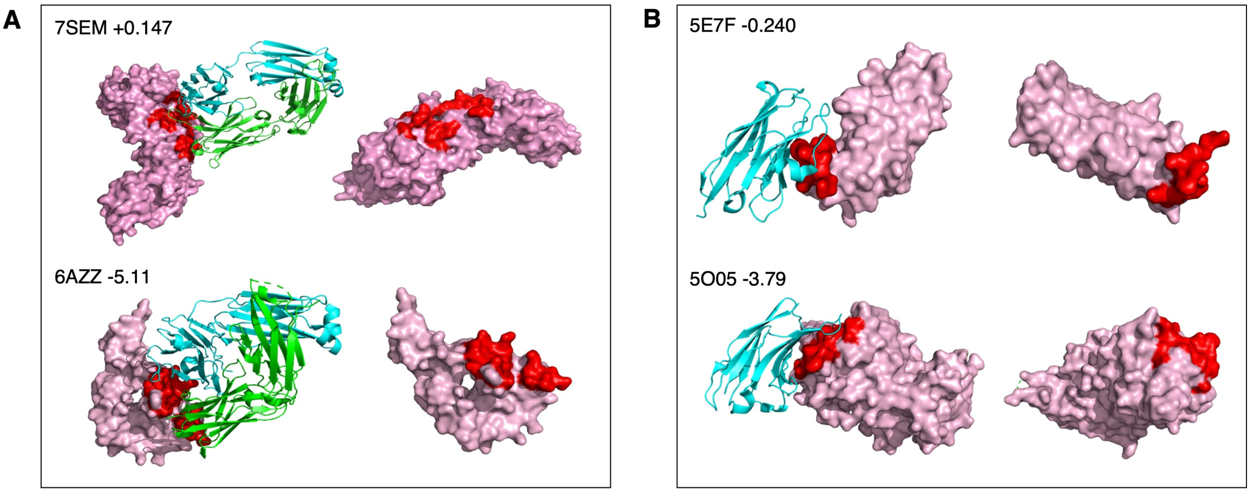


**Figure S13.** Visualising epitopes of structures alongside mean vertex defects values. The antibody chains are shown in blue and green ribbon, the antigen is shown as a surface representation and coloured pink and the epitope surface is shown in red. **(A)** The most convex (upper) and most concave (lower) epitopes targeted in Abs dataset. **(B)** The most convex (upper) and most concave (lower) epitope targeted in sdAbs dataset.

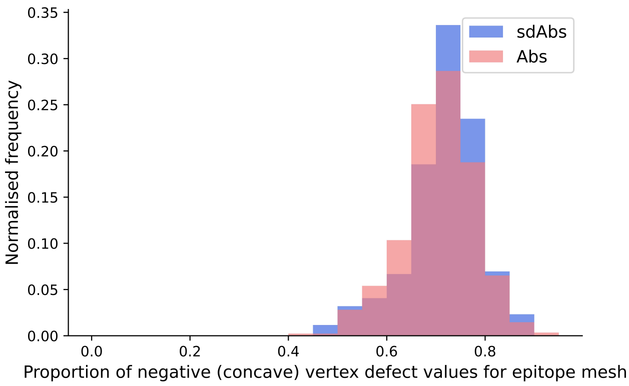


**Figure S14.** Distributions of the proportions of negative vertex defects values for interaction-defined epitopes of sdAbs (blue) and Abs (pink). The unpaired mean difference between sdAbs and Abs in the proportions of negative vertex defects values was -0.0089 (p-value = 0.15).

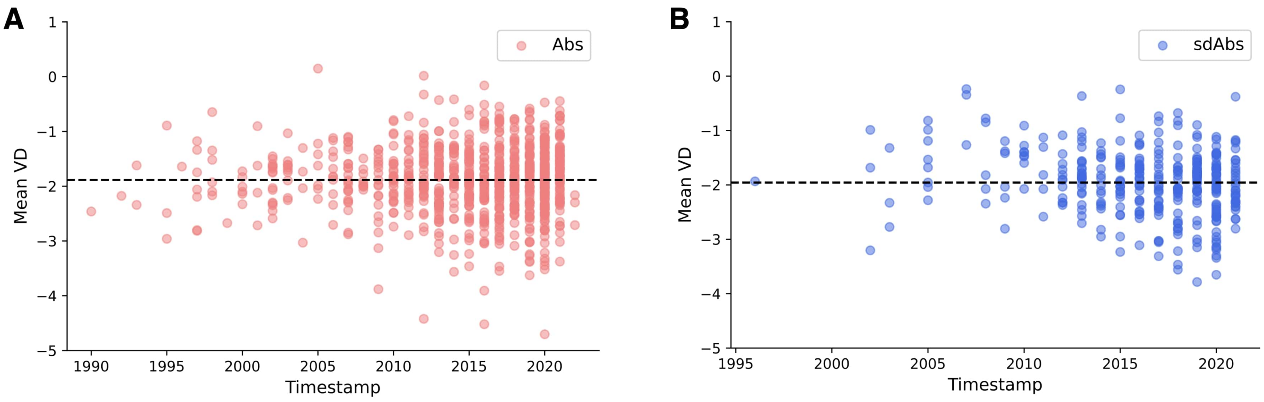


**Figure S15.** Distributions of mean vertex defects values for epitopes (for which more negative values indicate more concave epitopes) in our structural datasets, over time (PDB deposition date) indicate that the hypothesis that sdAbs target more concave epitopes does not come from older structures having more concave epitopes. The black dotted line indicates the mean mean vertex defects value of all structures in the dataset. **(A)** Mean vertex defects values over time for Abs dataset. **(B)** Mean vertex defects values over time for sdAbs dataset.

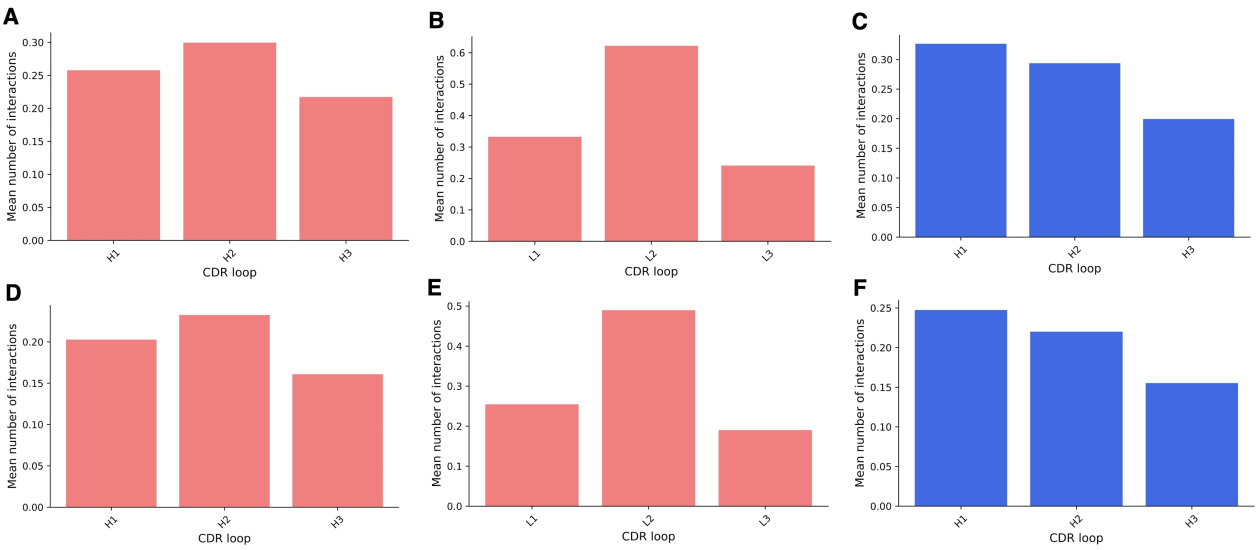


**Figure S16.** Bars show the mean number of interactions per CDR loop, normalised by loop length per complex. **(A), (B)** and **(C)** show these results for the distance-defined paratopes for the Abs VH, Abs VL and sdAbs respectively, whilst **(D), (E)** and **(F)** show the same but for the distance-defined paratopes.

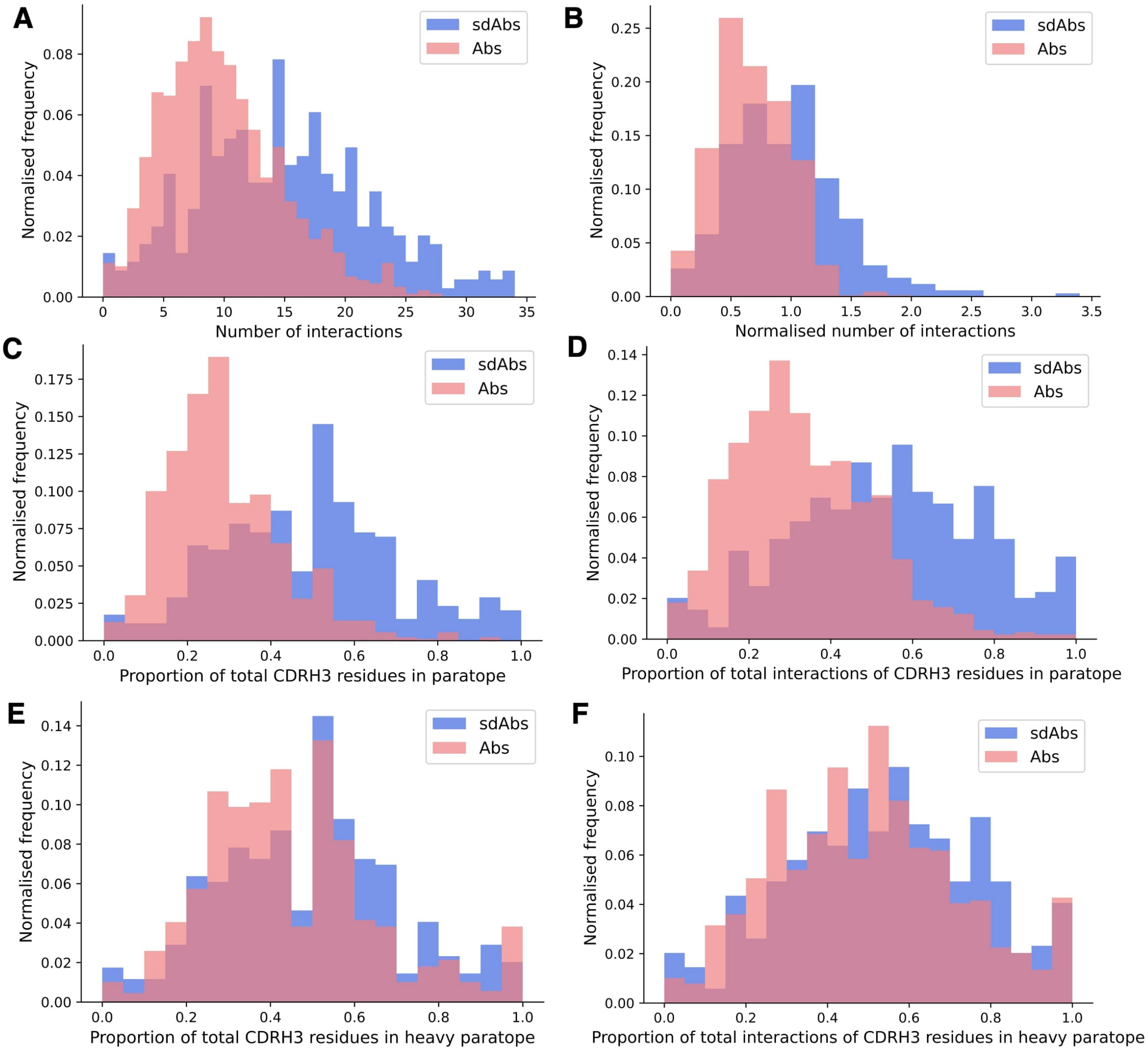


**Figure S17.** The CDR-H3 is of increased importance in sdAbs. **(A)** Distribution of the number of interactions established by CDR-H3 residues per complex in the sdAbs and Abs dataset (unpaired mean difference between sdAbs and Abs is -4.8, p-value « 0.01). **(B)** Normalised distribution for the number of interactions by the size (number of residues) of the CDR-H3 (unpaired mean difference between sdAbs and Abs is -0.27, p-value « 0.01). **(C)** Comparison of the fraction of number of residues contributed from the CDR-H3 loops for sdAbs and Abs. There is an unpaired mean difference of -0.19 between sdAbs and Abs with a p-value « 0.01. **(D)** Comparison of the fraction of number of interactions contributed from the CDR-H3 loops for sdAbs and Abs. There is an unpaired mean difference of -0.21 between sdAbs and Abs with a p-value « 0.01. **(E)** Comparison of the fraction of number of residues contributed from the CDR-H3 loops for sdAbs and Abs VH only. There is an unpaired mean difference of -0.075 between sdAbs and Abs VH with a p-value « 0.01. **(F)** Comparison of the fraction of number of interactions contributed from the CDR-H3 loops for sdAbs and Abs VH only. There is an unpaired mean difference of -0.052 between sdAbs and Abs with a p-value = 0.006.

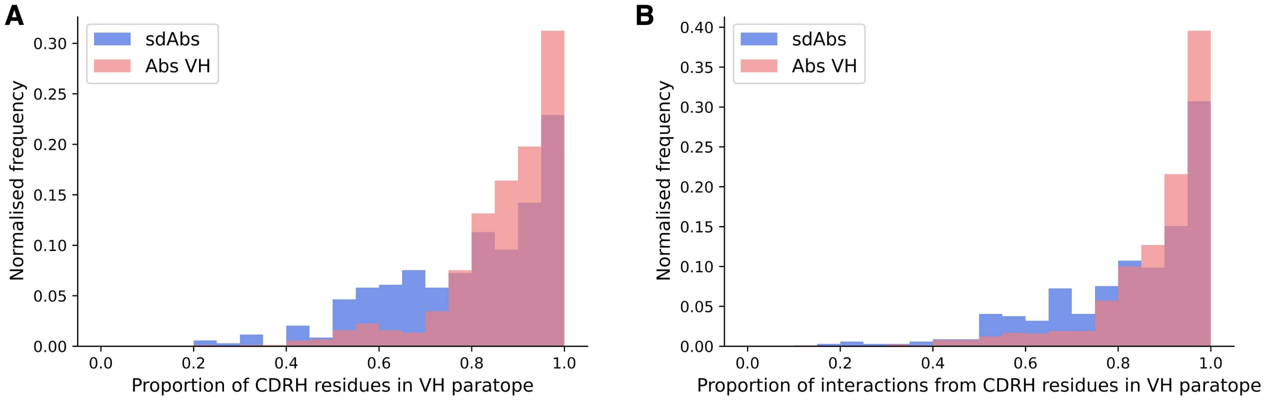


**Figure S18.** When comparing sdAbs paratope to the paratope residues from the Abs VH chains only, we observe that relatively fewer interactions are contributed by the CDRs in sdAbs, indicating that there is greater involvement of framework residues in the sdAbs paratope. **(A)** Comparison of the fraction of number of residues contributed from the CDR-H loops for sdAbs and Abs VH. There is an unpaired mean difference of 0.077 between sdAbs and Abs with a p-value « 0.01. **(B)** Comparison of the fraction of number of interactions contributed from the CDR-H loops for sdAbs and Abs VH. There is an unpaired mean difference of 0.055 between sdAbs and Abs with a p-value « 0.01.
